# LINE-associated cryptic splicing induces dsRNA-mediated interferon response and tumor immunity

**DOI:** 10.1101/2023.02.23.529804

**Authors:** Rong Zheng, Mikayla Dunlap, Jingyi Lyu, Carlos Gonzalez-Figueroa, Georg Bobkov, Samuel E. Harvey, Tracey W. Chan, Giovanni Quinones-Valdez, Mudra Choudhury, Amy Vuong, Ryan A. Flynn, Howard Y. Chang, Xinshu Xiao, Chonghui Cheng

## Abstract

RNA splicing plays a critical role in post-transcriptional gene regulation. Exponential expansion of intron length poses a challenge for accurate splicing. Little is known about how cells prevent inadvertent and often deleterious expression of intronic elements due to cryptic splicing. In this study, we identify hnRNPM as an essential RNA binding protein that suppresses cryptic splicing through binding to deep introns, preserving transcriptome integrity. Long interspersed nuclear elements (LINEs) harbor large amounts of pseudo splice sites in introns. hnRNPM preferentially binds at intronic LINEs and represses LINE-containing pseudo splice site usage for cryptic splicing. Remarkably, a subgroup of the cryptic exons can form long dsRNAs through base-pairing of inverted Alu transposable elements scattered in between LINEs and trigger interferon immune response, a well-known antiviral defense mechanism. Notably, these interferon-associated pathways are found to be upregulated in hnRNPM-deficient tumors, which also exhibit elevated immune cell infiltration. These findings unveil hnRNPM as a guardian of transcriptome integrity. Targeting hnRNPM in tumors may be used to trigger an inflammatory immune response thereby boosting cancer surveillance.

## Introduction

RNA splicing, a critical process that precisely removes introns from pre-mRNA, contributes remarkably to gene regulation and transcriptome diversity. In humans, introns constitute approximately 25% of the whole genome, representing an extensive expansion compared to lower eukaryotes and even other mammals.^1-3^ It has long been known that transposition by intronic retrotransposable elements is a major driver of intron expansion across various species.^4–6^ Through transposition, an abundance of repetitive DNA species, such as LINEs, SINEs, and ERVs, are inserted in introns of vertebrates, expanding intron length to several kilobases on average in human genes.^7, 8^

Long introns, however, contain copious amounts of pseudo splice sites that bear highly similar sequences to annotated splice sites in coding genes.^9–12^ Misusage of pseudo splice sites results in aberrant splicing, or cryptic splicing, which has been observed in several diseases.^13–16^ However, despite the presence of long introns in the human genome, splicing occurs in an accurate and precise manner. This suggests that a potent protective mechanism exists that represses cryptic splicing and ensures transcriptome integrity.

RNA-binding proteins (RBPs) are a class of proteins that regulate RNA processing, including splicing, localization, polyadenylation, and translation, through interaction with RNA and other proteins.^17–20^ Conceivably, RBP binding at pseudo splice sites may suppress cryptic splicing by masking these sites from recognition by the spliceosome. For example, hnRNPC was identified to suppress aberrant Alu exonization through preventing spurious U2AF65 binding at cryptic splice sites.^21^ While the full scope of cryptic splicing has been largely overlooked, expression of cryptic exons can be deleterious to normal gene expression, a mechanism that may partly account for the contribution of TDP-43 to neurodegenerative diseases such as amyotrophic lateral sclerosis (ALS).^22^ Moreover, MATR3 and PTBP1 were discovered to co-repress cryptic splicing in introns, and some of the cryptic exons may evolve into tissue-specific exons.^23^ Despite these findings, there is a lack of quantitative characterization of cryptic splicing regulated by RBPs, nor is there sufficient discussion of the clinical significance, especially in cancer, of RBP-regulated cryptic exons. Understanding the molecular mechanisms underlying pseudo splice site suppression can offer insights on the principles that ensure splicing-associated transcriptome integrity and potential therapeutics for mis-splicing related diseases.

In this study, we report the identification of hnRNPM as the top RBP that preferentially binds to introns. We observed a large quantity of cryptic exons expressed upon the loss of hnRNPM. hnRNPM binding is specifically enriched at intronic LINEs and is near splice sites of cryptic exons. A number of these LINE-bearing cryptic exons can form stem-like dsRNAs of roughly 300 base pairs in length, similar to dsRNA structures known to induce interferon responses.^24–26^ Intriguingly, the levels of hnRNPM expression and dsRNA-forming cryptic exons faithfully predict immune cell infiltration in tumors and thus patient survival outcomes. Together, our data provide a direct functional link between LINE-associated dsRNAs generated from hnRNPM-suppressed cryptic exons and interferon response induced tumor immunity.

## RESULTS

### hnRNPM preferentially binds to deep introns

The GENCODE v19 annotation of the human genome includes approximately 1.6 billion introns with a median length of 5 kB. These introns typically harbor many pseudo splice sites, sequences that resemble bona fide splice sites but are not used in normal splicing. Mechanisms that prevent splicing at pseudo splice sites remain incompletely understood. We hypothesized that a group of RBPs preferentially bind introns to repress pseudo splice sites, preventing recognition of such sites by the spliceosome, and thus protecting cells from generating cryptic exon-containing transcripts. To systematically survey RBPs that preferentially bind to introns, we analyzed RBP footprints using ENCODE eCLIP data of more than 150 curated RBPs.^27^ Comparing RBP binding peaks between the intronic and non-intronic genomic regions, we found that approximately 40% (67) of all examined RBPs had the majority of their binding sites in introns (Supplemental Table S1). 47 of those showed preferential intron-binding in both HepG2 and K562 cells. Among these, hnRNPM, MATR3, ILF3, PTBP1, and hnRNPL demonstrated the strongest preference for introns (**Fig. 1A**).

**Figure 1:**
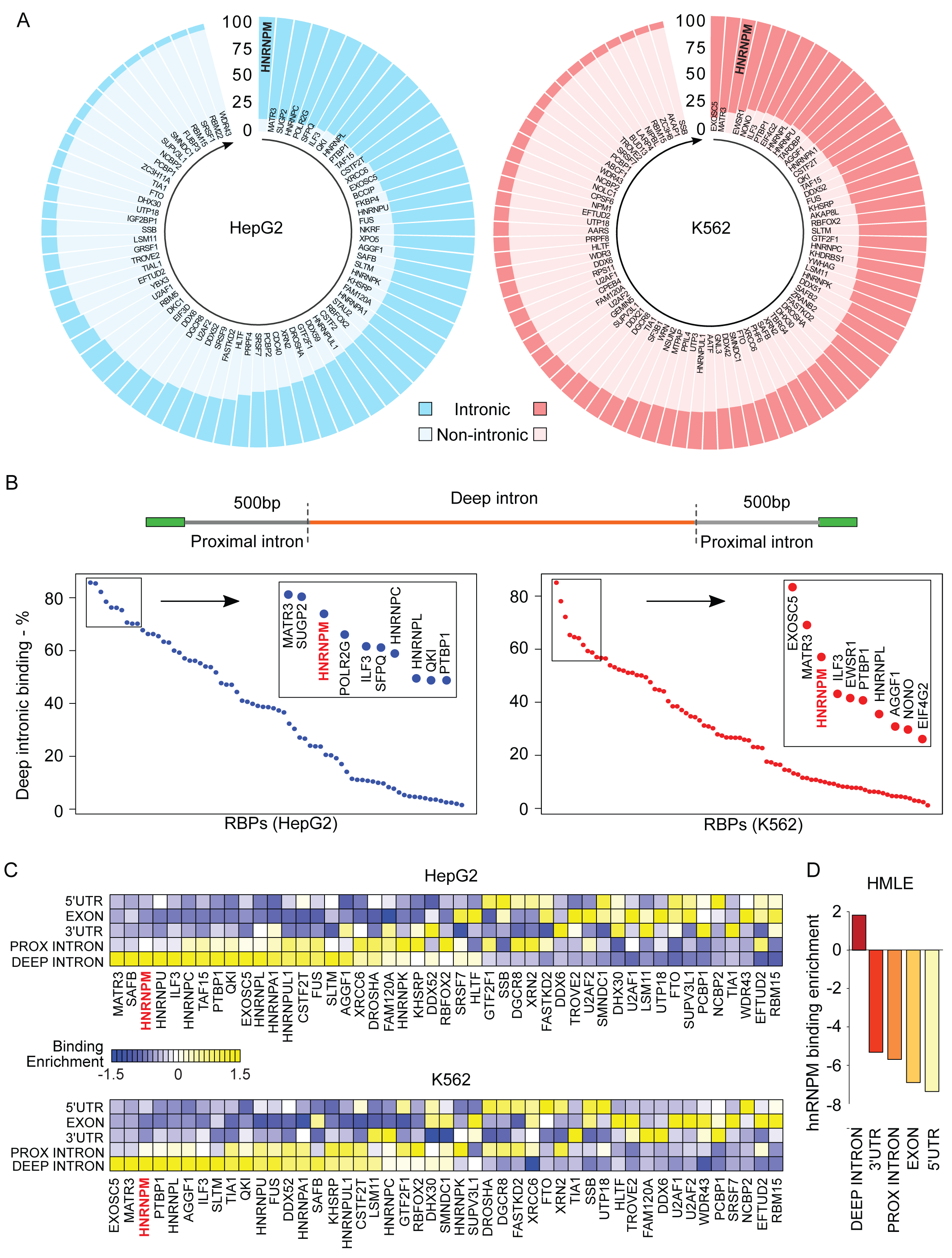
hnRNPM preferentially binds to deep introns. **A.** RBP binding distributions in intronic and non-intronic regions. Percent of intronic and non-intronic bindings of RBPs in HepG2 (n = 72) and K562 (n = 84) are shown in circular bar plots. Dark shades indicate intronic binding frequency, while light shades indicate their non-intronic binding frequency. RBPs that have less than 5% of their eCLIP binding peaks in introns are neglected. **B.** RBP binding distribution in deep introns. Upper diagram illustrates locations of deep and proximal introns. Dot plots in lower panel indicate RBP deep intronic binding frequency in HepG2 and K562 cell lines. Inserted box indicates top 10 ranked RBPs in each cell line. **C.** Heatmaps of RBP binding preferences to 5’UTR, exon, 3’UTR, proximal (PROX) intron, and deep intron across pre-mRNA. Binding preferences are shown for shared RBPs (n = 47) between HepG2 and K562 cells. **D.** hnRNPM iCLIP analysis in HMLE cells showing hnRNPM binding enrichment across pre-mRNA. Extended Data in Fig. 1 can be found in Supplemental Table S1.

Proximal intron regions, typically defined as up to 500 nucleotides adjacent to exons, are enriched with cis-regulatory elements which are bound by RBPs to regulate splicing. Intron regions that are beyond the proximal region are referred to as distal or deep introns (**Fig. 1B**, Top Panel), which have been overlooked in splicing regulation due to the assumption that regulatory sites are scarcely present in deep introns. Nonetheless, because the sizes of deep introns are often more than 5-fold longer than proximal introns, they harbor many pseudo splice sites that may be recognized by the splicing machinery under abnormal conditions, producing cryptic exons. We therefore segregated the intronic regions into proximal and deep introns and ranked the 67 RBPs identified in Fig. 1A based on their preference for binding at deep introns (**Fig. 1B**). Interestingly, the majority (43) of the 67 RBPs had more than 50% of their binding sites located in deep introns, suggesting that most RBPs that prefer intron-binding also predominantly occupy deep introns. hnRNPM, MATR3, ILF3, PTBP1 and hnRNPL continue to rank highest in their preferential binding to deep introns in both cell lines (See inset in **Fig. 1B**).

To account for length difference between proximal and deep introns, we assessed length-normalized binding distribution of the 47 shared RBPs between the two cell lines (Fig. 1A) and compared their binding preference in different regions, including 5’ UTR, coding exon, 3’UTR, proximal intron, and deep intron. The results demonstrated that MATR3, hnRNPM, ILF3, PTBP1, and EXOSC5 had a higher propensity to bind to deep introns, and this observation is highly reproducible in HepG2 and K562 cell lines (**Fig. 1C**). Meanwhile, we noted that a group of RBPs, such as RBM15, WDR43, and PCBP1, exhibited enriched binding to regions other than introns, suggesting that their normalized binding to non-intronic regions, i.e., 5’ UTR, exon, or 3’UTR, is greater than that in intronic regions.

Among the RBPs that preferentially bind to deep introns, hnRNPM captured our greatest interest since as much as 89% of its binding peaks reside in introns, placing it as the top RBP in HepG2 and the third highest ranking RBP in K562 cells (**Fig. 1A**). Moreover, hnRNPM was found to be an essential gene by CRISPR knockout screens,^28–30^ and its downregulation was reported to inhibit breast tumor metastasis.^31^ We experimentally examined hnRNPM binding by performing iCLIP in a human mammary epithelial cell line HMLE (**Fig. S1A**). We identified 54377 highly reliable hnRNPM binding sites at single nucleotide resolution (**Fig. S1B**). Approximately 90% of hnRNPM binding sites were in deep introns, 5% in proximal introns, and about 5% in UTRs and exons (**Fig. S1C**). De novo motif analysis of hnRNPM binding peaks revealed UG-rich motifs (**Fig. S1D**), highly similar to previously reported hnRNPM binding motifs,^32–34^ validating our iCLIP results. In addition, length-normalized binding distribution also showed that hnRNPM binding sites were most significantly enriched at deep introns (**Fig. 1D**). Collectively, by analyzing public ENCODE eCLIP datasets and iCLIP experiments in the HMLE cells, our results consistently showed that hnRNPM is a major RBP whose binding is significantly enriched in deep introns.

### Loss of hnRNPM results in global cryptic splicing

Because hnRNPM predominantly occupies deep introns, we hypothesized that hnRNPM binding to introns may mask pseudo splice sites from recognition by the splicing machinery. As a result, loss of hnRNPM could enable the use of pseudo splice sites, resulting in global cryptic splicing and transcriptome instability. To assess the impact of hnRNPM depletion on the cryptic splicing landscape, we conducted deep RNA-sequencing (RNA-seq) in control and hnRNPM knockdown (KD) HMLE cells (**Fig. S2A, S2B**). We developed a bioinformatic pipeline, CryEx, that detects all expressed exons including previously annotated exons or unannotated novel exons (**Fig. 2A**). We performed de novo assembly of RNA-seq reads to the genome using Stringtie and identified 493,975 expressed exons that are detected in at least half of the samples. These exons included 172,560 annotated and 321,415 unannotated exons. The majority of unannotated exons are located in deep introns.

**Figure 2:**
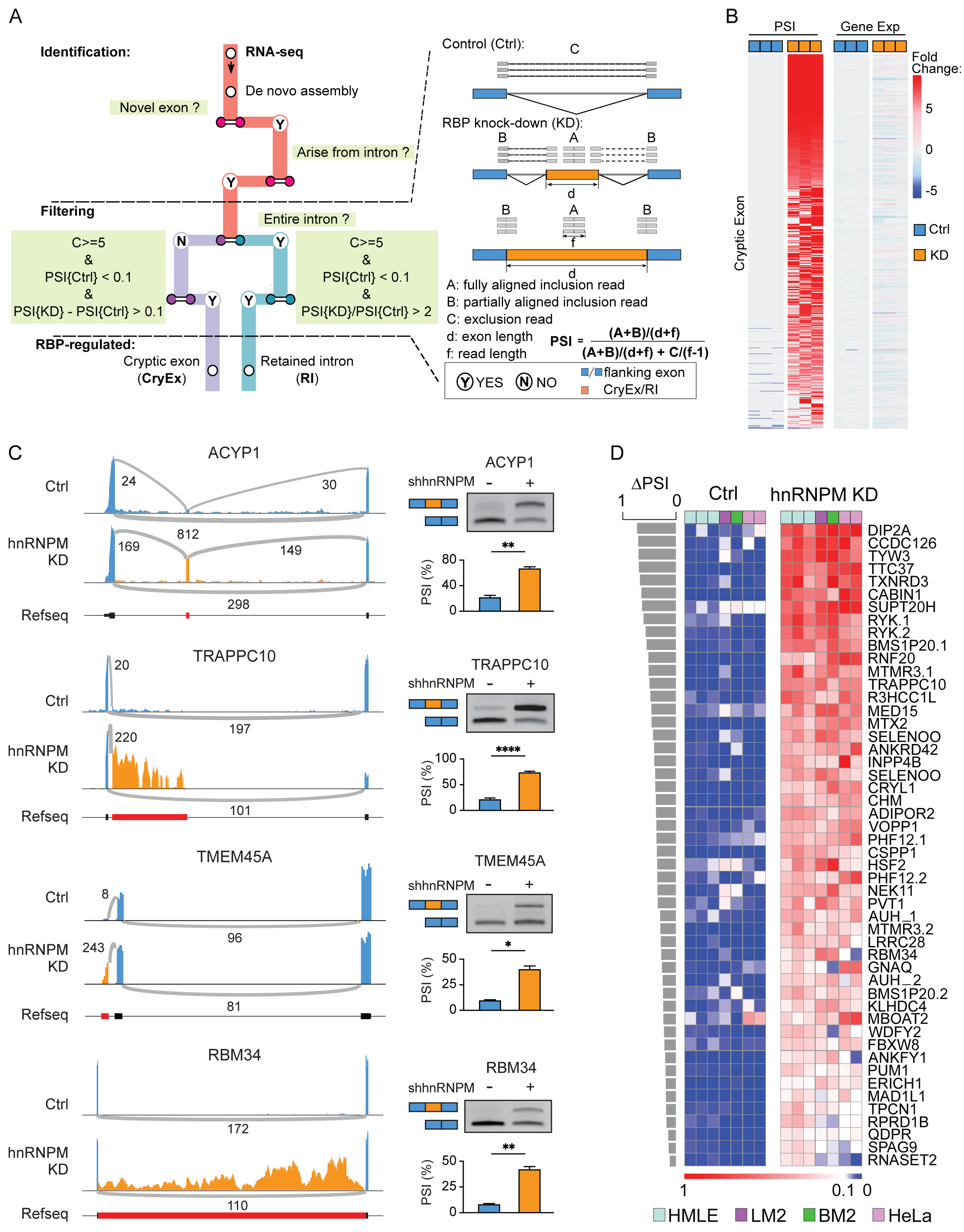
Loss of hnRNPM results in global cryptic splicing. **A.** A workflow to identify cryptic splicing from RNA-seq data. Showing inside dashed lines are the criteria to report a cryptic exon event, including number of reads at the exclusion junction (indicated by C) and a percent spliced in (PSI) metric calculated from A, B, C, d and f. **B.** Heatmaps showing cryptic exon production is independent of alterations of gene expression changes. *Left*, heatmaps of PSI fold change in control (Ctrl, blue) and hnRNPM knockdown (KD, Orange) cells. *Right*, fold changes of transcript levels of genes that contain cryptic exons in hnRNPM KD cells compared to control cells. **C.** Representative examples of hnRNPM-repressed cryptic exons. *Left*, sashimi plots of cryptic splicing events. Height of the peaks represent number of reads aligned, and the thickness of arches represent numbers of spliced reads. *Right*, semi-qPCR validations. Agarose gel images are shown on the top. Quantifications of the cryptic splicing products represented as PSI values are shown at the bottom. **D.** Heatmaps of PSI values of top 50 representative hnRNPM-repressed cryptic exons identified from HMLE cells in four cell lines with hnRNPM KD (HMLE, LM2, BM2, HeLa). Cryptic events are ranked from high to low based on ΔPSI values (ΔPSI = PSI{KD} – PSI{Ctrl}). Extended Data in Fig. 2D can be found in Supplemental Table S2.

We used differences of Percent-Spliced-In (ΔPSI) to quantify splicing changes between control and hnRNPM KD conditions. CryEx analysis of annotated exons in control and hnRNPM KD cells revealed 716 alternatively skipped exons (ASEs) with an absolute ΔPSI greater than 0.1. These alterations are highly consistent with results from another splicing pipeline rMATs,^35^ with a significant correlation co-efficient of 0.9, demonstrating the accuracy of our CryEx tool in identifying alternatively spliced exons and quantifying their inclusion rates (**Fig. S2C**). Furthermore, the majority of ASEs in hnRNPM KD cells are inclusion events, which is consistent with previous findings that hnRNPM promotes exon skipping ^36^ (**Fig. S2D**).

Next, we proceeded to analyze novel splicing events identified by CryEx. We found more than 1000 novel splicing events that occur due to the loss of hnRNPM that are essentially absent in control cells. Most of the genes containing cryptic exons do not show differential expression of their mRNA transcripts (**Fig. 2B, S2E**), indicating that the observed cryptic events are generated by abnormal splicing rather than changes in transcription. Experimental validation by semi qRT-PCR of nearly 80 randomly selected ASEs and cryptic splicing events showed a Pearson correlation coefficient above 0.9 between experimental and RNA-seq quantifications, further supporting the accuracy of our CryEx pipeline (**Fig. S2F, S2G**).

Most (78%) cryptic exons reside in annotated introns, between the upstream and downstream annotated exons, resembling a canonical exon (**Fig. 2C**, ACYP1). Among these cryptic exons, 68% do not preserve the reading frame (length being not a multiple of three). Consequently, their inclusion may cause a frameshift in the mRNA transcript and/or generate stop codons that could render the altered RNAs to become targets of nonsense mediated decay (NMD) (**Fig. S2H**). The second group of cryptic splicing events, exemplified by TRAPPC10 (**Fig. 2C**), contains a stop codon and a polyA signal downstream of the cryptic stop codon, generating a truncated mRNA with a novel 3’UTR (**Fig. S2I**). In this type of cryptic splicing, we detected only splice junction reads between the 5’ splice site of the annotated exon and the 3’ splice site of the cryptic exon, but not between the 5’ splice site of the cryptic exon and 3’ splice site of its downstream exon, which further reinforced our prediction on the production of a novel 3’UTR. The third type of cryptic splicing occurs upstream of the first exon. This extends the 5’UTR and potentially causes the usage of a new translation start site upstream. As expected, only the 5’ splice site of the cryptic exon is detected in the splice junction reads (**Fig. 2C**, TMEM45A). Additionally, cryptic splicing can also cause intron retention. These retained introns have an average size of approximately 7 kB. Many of them, such one in RBM34, contain a stop codon in the retained intron, potentially producing a truncated protein. Gene ontology analysis of the cryptic spliced genes shows that they fall into a broad range of biological pathways (**Fig. S2J**), suggesting a global impact of cryptic splicing upon the loss of hnRNPM.

To directly examine the role of hnRNPM in repressing cryptic splicing, we constructed a splicing reporter minigene that contains the cryptic exon of the ADIPOR2 gene, and its adjacent introns flanked by two constitutive exons (**Fig. S2K**, Upper). Splicing of the ADIPOR2 cryptic exon was gauged using semi-qRT-PCR to measure the percentage of cryptic exon inclusion (**Fig. S2K**, Middle). Co-transfection with the minigene construct and different amount of hnRNPM cDNA in 293FT cells demonstrated that hnRNPM represses ADIPOR2 cryptic exon inclusion in a dose-dependent manner, resulting in a decrease of cryptic exon inclusion from 92% in control cells to 69% in cells transfected with 800 ng of hnRNPM cDNA. Moreover, we examined the impact of hnRNPM on repressing cryptic retained introns. We inserted the sequence of intron that is retained in MED15 into the splicing reporter minigene (**Fig. S2K**, Upper) and found that hnRNPM also represses inclusion of the MED15 intron in a dose-dependent manner (**Fig. S2K**, Lower). These results indicate that hnRNPM directly represses cryptic splicing.

An intriguing observation is that cryptic splicing does not occur at random. Many hnRNPM-repressed cryptic exons identified in HMLE cells are highly replicable across cell lines. As shown in **Fig. 2D**, the top 50 hnRNPM-repressed cryptic exons are also consistently detected in the lung metastatic breast cancer line LM2, bone metastatic breast cancer line BM2, and the cervical cancer line HeLa upon the loss of hnRNPM. The recurring non-random cryptic splicing detected in multiple cell lines suggests that the pseudo splice sites are repressed in normal conditions and become recognized by the splicing machinery when hnRNPM levels are low or depleted.

### hnRNPM binding overlaps with cryptic splice sites at deep intronic LINEs

To delineate the mechanism of hnRNPM-mediated cryptic exon repression, we systematically examined the sequence characteristics of cryptic exons and pseudo splice sites bound by hnRNPM using iCLIP-seq. First, we found that hnRNPM-repressed cryptic exons are derived from introns (and genes) that are longer than those of hnRNPM-mediated annotated ASEs or annotated background exons (and their associated genes) (**Fig. 3A**, **S3A**). Second, as high as 35% of hnRNPM-repressed cryptic splicing events are hnRNPM-bound, significantly higher than that of hnRNPM-regulated and annotated alternatively spliced exons, where only 5% had detectable hnRNPM binding close by (**Fig. 3B**). Third, roughly 95.6% of the hnRNPM-bound cryptic exons are in deep introns, while merely 4.39% being embedded in proximal introns (**Fig. 3C**). The average distance between the cryptic exons and the nearest annotated exons is roughly 8 kB, further supporting that hnRNPM-bound cryptic exons are enriched in deep introns (**Fig. S3B**).

**Figure 3:**
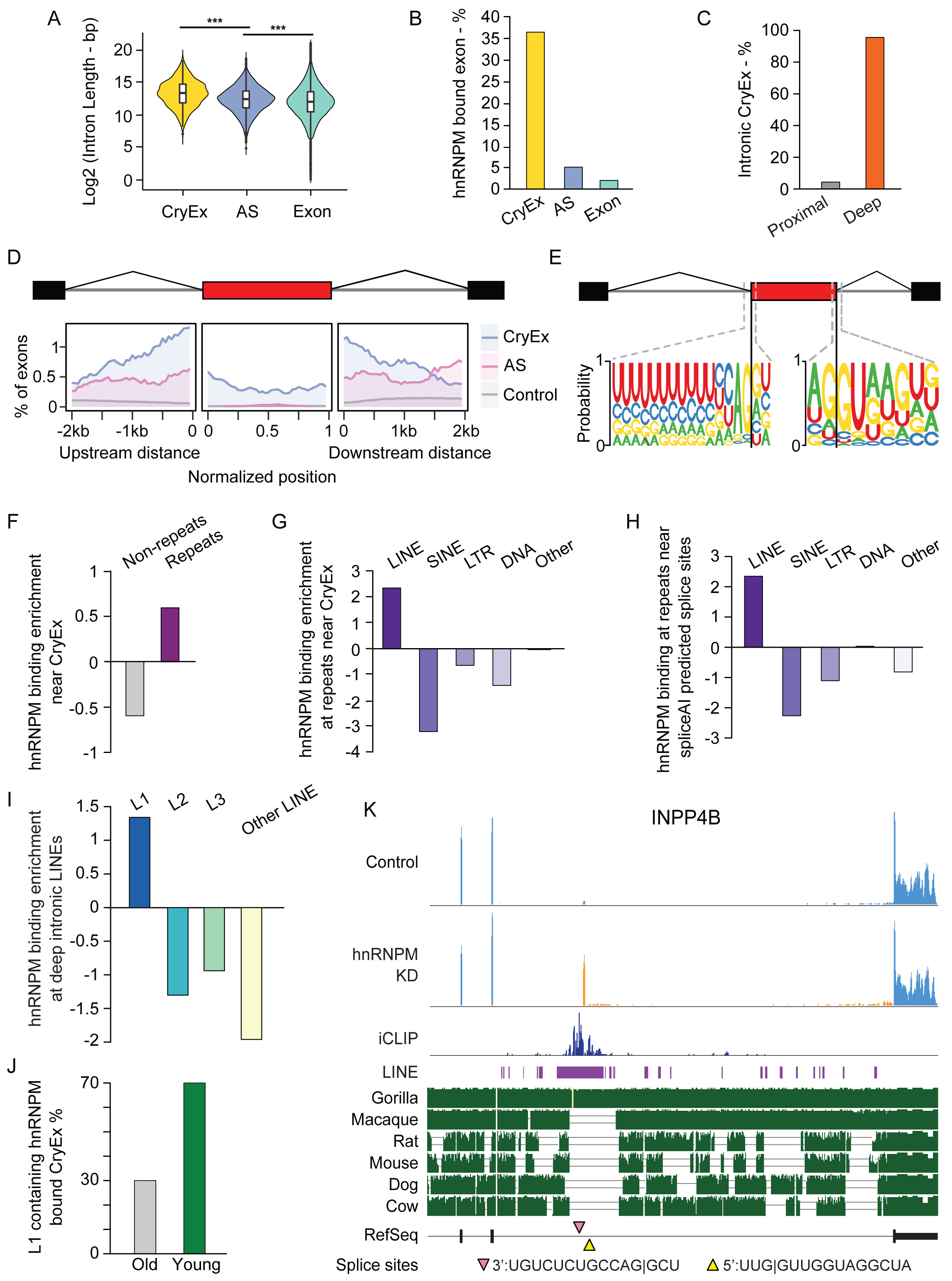
hnRNPM binding overlaps with cryptic splice sites at deep intronic LINEs. **A.** A violin plot showing length distribution of introns from hnRNPM-regulated cryptic exons (CryEx, Yellow), alternative spliced exons (AS, Blue), and annotated background exons. **B.** A bar plot showing percentage of hnRNPM binding to hnRNPM-regulated cryptic exons (CryEx, Yellow), alternative spliced exons, and annotated background exons. **C.** A bar plot showing that most intronic cryptic exons reside in deep introns. **D.** The metadata profile showing that hnRNPM is most significantly enriched at the immediate upstream and downstream of its repressed cryptic exons compared to its binding to alternatively spliced exons or all other annotated exons. hnRNPM binding in a +/ -2kb window flanking three groups of exons are shown. **E.** Consensus motifs at splice sites of hnRNPM-repressed cryptic exon. **F.** A bar plot showing the enrichment of hnRNPM binding at repeat elements near cryptic exons within deep introns. **G.** A bar plot showing that hnRNPM binding near cryptic exons is specifically enriched at LINEs. **H.** A bar plot showing hnRNPM binding at spliceAI predicted splice sites is most enriched at LINEs. **I.** A bar plot showing that hnRNPM binding is most enriched at L1. **J.** A bar plot showing that evolutionarily young L1s are the major L1 found in hnRNPM bound and repressed cryptic exons. **K.** An example of an hnRNPM-repressed cryptic exon in the INPP4B gene. Top two IGV tracks represent RNA-seq reads aligned in control and hnRNPM KD cells. The third track represents hnRNPM iCLIP binding. LINE elements are depicted as purple boxes under iCLIP peaks, and the LINE that this cryptic exon reside in is L1. A snapshot of the Primates Multiz Alignment & Conservation tracks on UCSC genome browser is shown as green tracks that includes 6 species, top 2 of which are primates. The 3’ (pink) and 5’ (yellow) splice sites of the cryptic exon are shown as triangles on the RefSeq, and they as well were predicted by SpliceAI. Sequences of the splice sites were included at the bottom of the panel.

To evaluate hnRNPM binding intensity across all human introns and exons, we conducted a metagene analysis of iCLIP data, which revealed significantly more frequent hnRNPM binding around its regulated exons, including both cryptic and alternative exons, compared to the background (**Fig. 3D**). While hnRNPM showed no binding preference at 5’ or 3’ splice sites, it had a significantly higher binding percentage near its repressed cryptic exons compared to annotated alternative exons. The motif derived from hnRNPM iCLIP binding near cryptic exons is GU-rich (**Fig. S3C**), similar to hnRNPM’s binding motif on global RNA (**Fig. S1D**) and from previous reports.^36, 37^ Using reported hnRNPM motifs for hnRNPM distribution further confirmed the results of iCLIP metagene analysis (**Fig. S3D**). Additionally, using RNA oligos derived from iCLIP hnRNPM peaks near the cryptic exons of ADIPOR2 and TRAPPC10, we validated hnRNPM binding by RNA pull-down (**Fig. S3E**). These results show that hnRNPM is enriched in introns immediately next to cryptic exons.

We then asked whether the cryptic exons repressed by hnRNPM contain the same consensus motifs at 5’ or 3’ splice sites as annotated exons. Indeed, both hnRNPM-repressed cryptic exons and annotated alternative exons regulated by hnRNPM share nearly identical 5’ and 3’ splice site motifs, which are indistinguishable from the reported conserved 5’ and 3’ splice site motifs (**Fig. 3E, S3F**).^38^ These results provide additional evidence that hnRNPM-repressed cryptic exons contain conserved splice sites, further supporting the model that hnRNPM binding represses pseudo splice site recognition in deep introns and thus prevents cryptic splicing. The conserved pseudo splice sites repressed by hnRNPM also provide an explanation to our observations that hnRNPM-repressed cryptic splicing does not occur at random (**Fig. 2D**).

Next, we sought to understand the sequence elements preferentially bound by hnRNPM. We first divided deep introns into repeat and non-repeat groups based on RepeatMasker annotations^39, 40^ and examined hnRNPM binding enrichment near cryptic exons located in these two groups. Interestingly, hnRNPM binding is distinctively enriched at cryptic exons in repeat regions (**Fig. 3F**). Further classifying the repeats based on their repetitive DNA distribution and sequence complexity showed that hnRNPM is most specifically enriched at long interspersed nuclear elements (LINEs) in deep introns compared to other repeat types (**Fig. 3G**).

Since our results suggest that hnRNPM binding represses pseudo splice site usage, we applied SpliceAI,^41^ a computational tool that uses artificial intelligence to predict splice sites, to predict potential splice sites in deep intronic repeats. We found that SpliceAI-predicted splice sites are particularly highly enriched for hnRNPM binding sites at LINEs (**Fig. 3H**), supporting our findings that hnRNPM binding shields pseudo splice sites that are highly enriched in LINEs from recognition. Furthermore, normalizing hnRNPM binding sites to the length of different LINE elements showed that hnRNPM binding is most enriched in LINE1 (L1), the most abundant LINE in humans (**Fig. 3I**).

As the most abundant non-long terminal repeat retrotransposons found in the genome, L1 insertion is a key contributor to intron expansion during evolution.^42^ We therefore assessed if evolutionary age of human L1 impacts cryptic exons repression by hnRNPM. Among all hnRNPM binding peaks associated with cryptic exons that contain L1, we found 70% belong to young L1s that are conserved only in primates, while the remaining 30% fall into the category of evolutionarily old L1s (**Fig. 3J**). Shown in **Fig. 3K** is a representative example of INPP4B, where hnRNPM binds near its cryptic exon. hnRNPM binding sites fall on an L1 element conserved between human and the closely related gorilla genomes, but not in other species. This preferential binding of hnRNPM to evolutionarily young L1s supports a role for hnRNPM in repressing cryptic exons that are embedded in young L1s. It may also suggest that hnRNPM plays a critical role to repress cryptic splicing in intron expansion during evolution.

### hnRNPM-repressed cryptic exons form cytoplasmic long stem-like dsRNAs

Since repetitive sequences are prone to forming secondary structures, we speculated that long cryptic exons repressed by hnRNPM may have a high tendency to form secondary structures. Surveying several long cryptic exons for potential RNA structures using RNAFold^43^ led us to a surprising finding that they are predicted to form remarkably long and stable dsRNAs (**Fig. 4A, S4A**). In addition to containing the LINE elements (purple in **Fig. 4A**), these dsRNA regions also contain ALU elements, a primate-specific class of short interspersed elements (SINEs).^44^ The nearly perfect base pairing inside these dsRNAs is formed from inverted repeats of Alu elements, shown as red blocks that are scattered between LINE elements (**Fig. 4A, S4A**). These dsRNAs can reach near 300 base pairs long and show drastically decreased Gibbs free energy (ΔG), suggesting structure stability.

**Figure 4:**
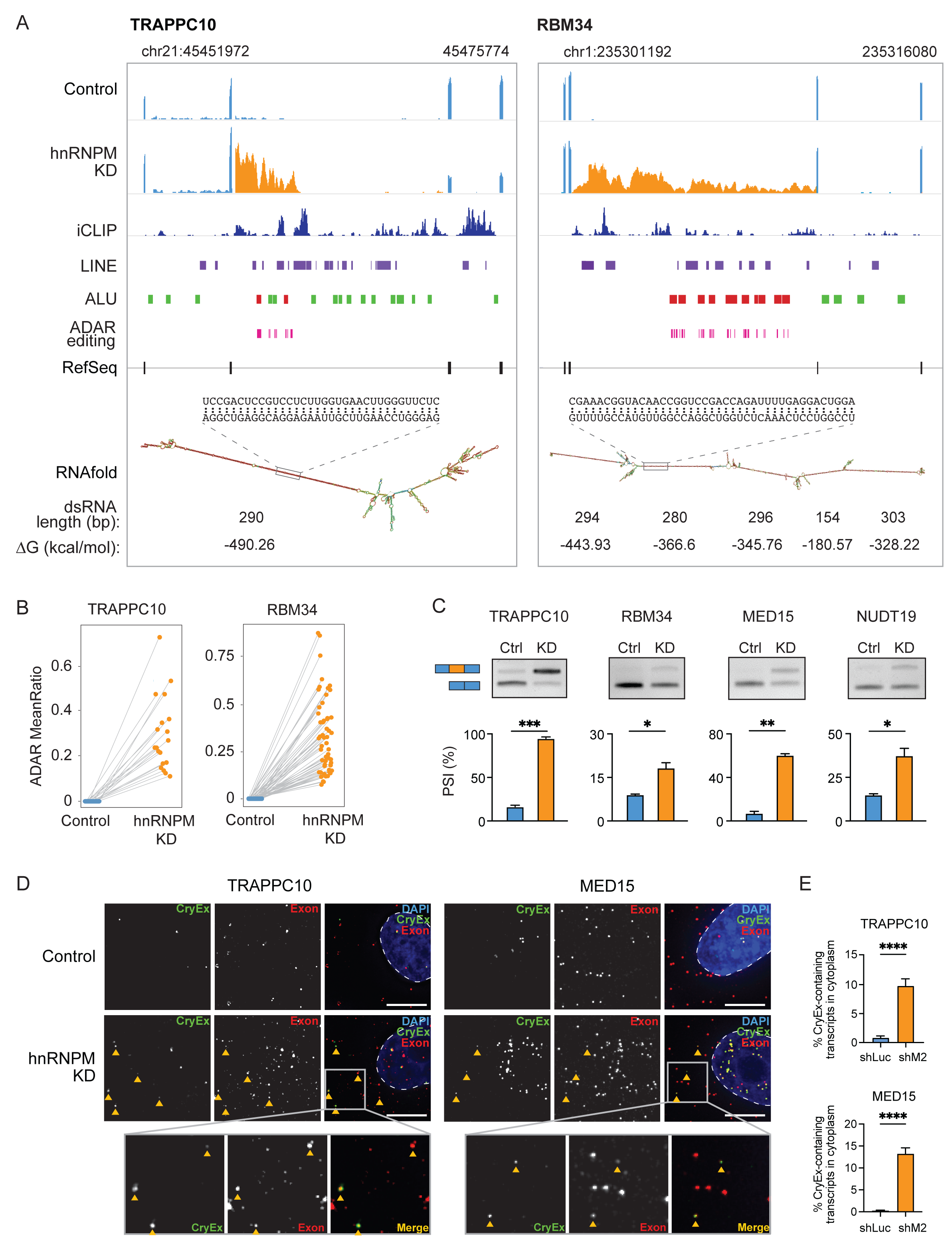
hnRNPM-repressed cryptic exons form cytoplasmic long stem-like dsRNAs. **A.** Representative examples of dsRNA-forming cryptic exons. *Left,* TRAPPC10. *Right*, RBM34. Top two IGV tracks indicate number of reads aligned. Third track represents hnRNPM iCLIP binding data. Purple boxes under each iCLIP track are LINE elements. Green boxes are Alu elements. Red boxes are Alu elements that are predicted to form dsRNAs via RNAfold. Pink boxes are ADAR editing sites. Seventh track indicates RefSeq annotation. Eighth track representing secondary structure of cryptic exon predicted by RNAfold. Approximate lengths of highly base-paired dsRNA regions are indicated beneath the structures together with predicted folding free energies (ΔG). A fragment of the dsRNA region is indicated above the secondary structure. **B.** Dot plots showing ADAR editing in predicted dsRNA-forming regions. **C.** semi-qRT-PCR using cytoplasmic mRNA showing a significant increase of dsRNA-forming cryptic exons in hnRNPM KD cells. **D.** RNA fluorescence in situ hybridization (RNA-FISH) images of dsRNA containing cryptic exons (CryEx, Green) and annotated exons (Exon, Red) for TRAPPC10 and MED15 genes. Nuclei are stained with DAPI, and nuclei boundaries are shown as dotted lines. Arrows indicate overlapped dsRNA-containing cryptic exon and annotated exon foci in the cytoplasm. Scale bars, 10 µm. **E.** Quantification of cytoplasmic dsRNA-forming cryptic exon-containing mRNA in control and hnRNPM KD cells (mean +/ -SEM, n = 3, 8 - 25 cells per experiment, two-tailed unpaired student’s t test).

dsRNAs are the major substrates of ADARs, which edit adenosine to inosine by deamination.^45, 46^ To examine whether the predicted dsRNAs exist in cells, we analyzed ADAR-mediated editing sites on the cryptic exons with predicted dsRNA structures in control and hnRNPM KD cells. If cryptic exons form dsRNAs, they will be subjected to ADAR editing, therefore accumulating A-I editing sites. We analyzed A-I editing of four cryptic exons that were predicted to form long stem-like dsRNA structures and showed larger ΔPSIs. Strikingly, in all four examined cryptic exons residing in TRAPPC10, RBM34, NUDT19, and MED15, we observed a drastic increase of both ADAR editing sites and editing levels within predicted dsRNA regions in hnRNPM KD cells (**Fig. 4B, S4B**).

To further examine whether the cryptic exons contain sequences that can form double-stranded hairpins, we used cDNA reversely transcribed from RNA of the hnRNPM KD cells and amplified the predicted dsRNA regions in the TRAPPC10 cryptic exon by high fidelity PCR. DNA sequencing of the PCR product showed virtually perfect base-pairing of the inverted repeats (**Fig. S4C**). These results support the idea that hnRNPM-repressed cryptic exons can form dsRNAs.

To further investigate whether dsRNA-containing transcripts exist in the cytoplasm, we isolated the cytoplasmic fraction of HMLE cells and measured cryptic exon inclusion by semi qRT-PCR (**Fig. 4C, S4D**). As expected, control cells predominantly express normally spliced transcripts with little or no detectable inclusion of cryptic exons. However, cryptic transcripts are readily detected in the cytoplasm of hnRNPM KD cells, and their levels are significantly higher compared to controls, proving evidence that dsRNA-containing cryptic transcripts exist in the cytoplasm (**Fig. 4C**). The existence of dsRNA-containing transcripts in the cytoplasm was further confirmed using the cell line LM2 (**Fig. S4E**). As a side note, although nearly no MED15 cryptic exon in control cells was detected (**Fig. 4C, S4E**), we observed a 50% inclusion of this exon when using whole cell lysates (**Fig. S2G**). This inclusion level, although markedly lower than that of 82% in hnRNPM KD cells, is considerably higher than the inclusion level in the cytoplasmic fraction. We speculate that a large fraction of MED15 cryptic transcripts, although produced, are trapped in the nucleus in control cells and are not exported to the cytoplasm. This is in line with the observed ADAR editing sites in control cells, where RNA from whole cell lysates was used. This result also suggests the existence of a surveillance mechanism in control cells, where intron-containing transcripts are prevented from been exported and translated to aberrant proteins.

To directly visualize the presence of dsRNA-containing cryptic transcripts in the cytoplasm, we performed RNA fluorescence in situ hybridization (RNA-FISH) using RNA probes that span the dsRNA region and its proximal cryptic exon regions (Green color, denoted as CryEx) and RNA probes that span annotated exons in the mRNA transcripts (Red color, denoted as Exon). Consistent with semi qRT-PCR results in Fig. 4C, almost no dsRNA-containing cryptic exon signals were detected in the cytoplasm of control cells but were prominently found in that of hnRNPM KD cells (**Fig. 4D**). Quantification results show that 9.76% and 12.69% of cytoplasmic TRAPPC10 and MED15 transcripts contain the dsRNA cryptic exons, respectively (Fig. 4E). Collectively, these multiple lines of evidence indicate that hnRNPM-repressed cryptic exons produce cytoplasmic dsRNAs.

### Cryptic splicing produced dsRNA elevates type-I interferon response

Mounting evidence have demonstrated that dsRNAs of more than 30-bp in length act as a potent trigger of antiviral immunity.^47–49^ Host dsRNA sensors, including DDX58 (RIG-I), IFIH1(MDA5), and TLR3, recognize these dsRNAs and trigger a cascade of pathways to activate type I interferon (IFN-I) response and upregulate overall expression of interferon-stimulated genes (ISGs) as a cellular response against viral infection.^25, 50^ Because our results suggested that cytoplasmic dsRNAs are produced by cryptic splicing in hnRNPM KD cells, we hypothesized that these dsRNAs act as a viral mimic, which are recognized through cellular dsRNA sensing. This subsequently results in activation of the IFN-I response and increased ISG production (**Fig. 5A**).

**Figure 5:**
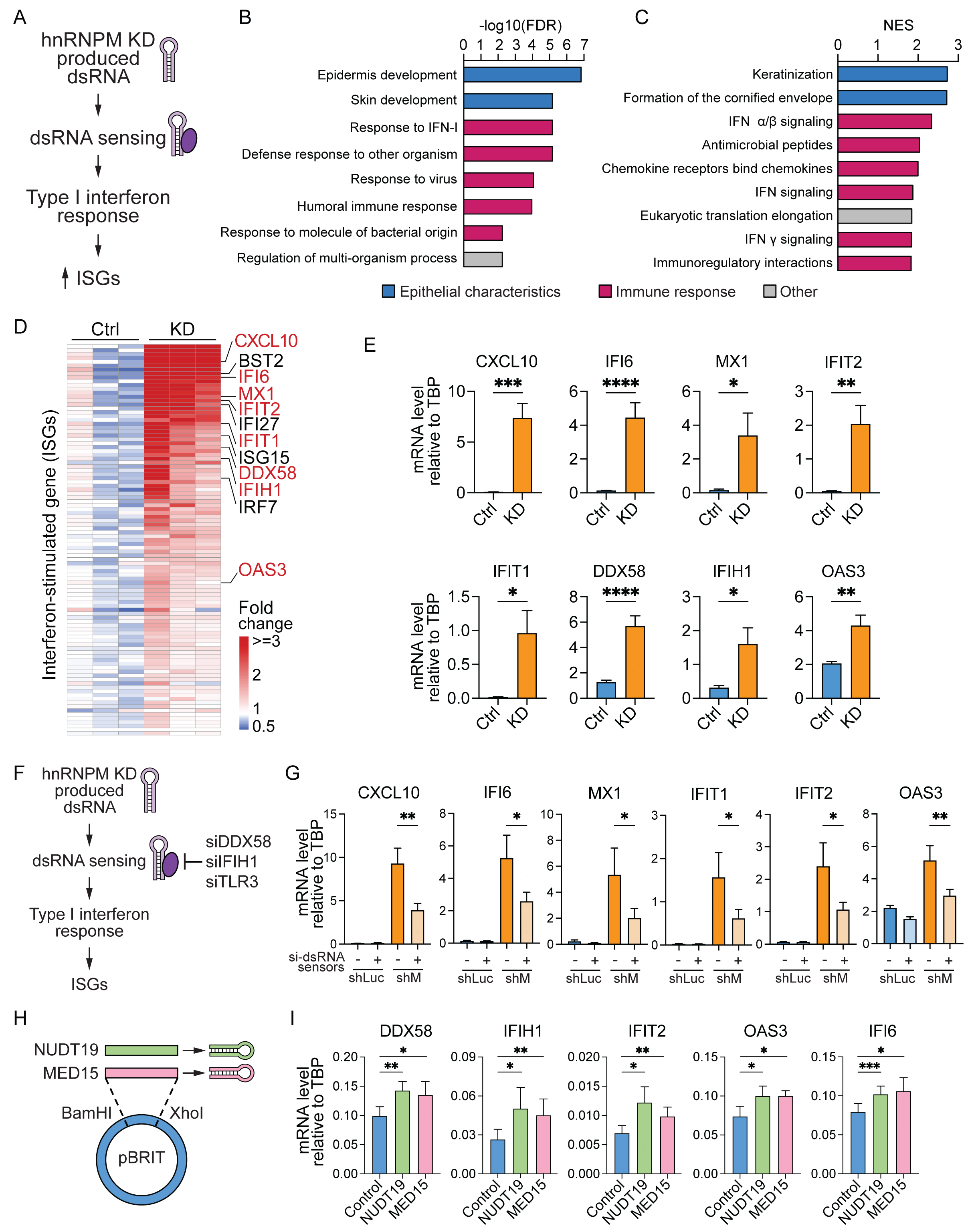
Cryptic splicing-produced dsRNA elevates type-I interferon response. **A.** A model of hnRNPM KD induced type I interferon response through the production of dsRNA. **B,C.** Gene Ontology and Gene Set Enrichment Analysis showing top biological processes and pathways positively enriched in hnRNPM KD cells. Immune-related GO terms and pathways are indicated in red. Epithelial characteristics related gene sets are shown in blue, and other terms are shown in grey. **D.** A heatmap of ISG expression showing that ISGs are globally upregulated in hnRNPM KD cells. Relative expression was calculated as mean FPKM fold change versus control. ISGs involved with virus stimulated interferon response are highlighted in red. **E.** qRT-PCR analyses on ISGs highlighted in **D**. Data is relative to TBP (mean +/ - SEM, n=10 biological replicates, two-tailed paired student’s test). **F.** A model showing how siRNA treatment of dsRNA sensors (DDX58, IFIH1, TLR3) may affect downstream activation of type I interferon response and ISG expression. **G.** qRT-PCR analyses showing that siRNA knockdown of dsRNA sensors in hnRNPM KD cells reduces ISGs expression significantly. Comparison of bars 1 and 3 shows upregulation of ISGs in hnRNPM KD cells. Comparison of bars 4 and 3 shows downregulation of ISGs upon siRNA depletion of dsRNA sensors in hnRNPM KD cells. Data is relative to TBP (mean +/ - SEM, n=5 biological replicates, two-tailed paired student’s test). **H.** A diagram showing the plasmid design that expresses dsRNA forming cryptic exon of NUDT19 or MED15. **I.** qRT-PCR analyses of ISGs expression in control, NUDT19- or MED15-cryptic exon expressing cells. Ectopic expression of dsRNA-forming cryptic exons significantly upregulated ISG levels. Data is relative to TBP (mean +/ - SEM, n=3 biological replicates, two-tailed paired student’s test).

We compared transcriptomes in control and hnRNPM KD cells. Gene ontology (GO) analysis (**Fig. 5B**) and gene set enrichment analysis (GSEA, **Fig. 5C**) showed that several antiviral signaling pathways, including IFN-I responses, were among the most significantly positively enriched pathways in hnRNPM KD cells. Additionally, terms associated with the epithelial cell state were also highly positively enriched in hnRNPM KD cells. This is consistent with previous reports that hnRNPM plays an important role in promoting epithelial-mesenchymal transition (EMT).^31^ Previous studies have also shown that hnRNPM deficiency impairs cell fitness.^33, 51^ Consistent with this finding, we found that cell-cycle pathways are significantly downregulated in response to hnRNPM knockdown (**Fig. S5A**).

Next, we curated a signature of ISG regulation that incorporates dsRNA-specific antiviral genes^52^ and hallmark interferon α, β, and γ gene sets.^53^ As predicted, hnRNPM KD cells showed significant upregulation of a broad spectrum of ISGs (**Fig. 5D**). The top 40 upregulated ISGs exhibited greater than 4-fold increase in hnRNPM KD cells (**Fig. S5B**). These ISGs include the dsRNA sensors DDX58 and IFIH1 and the cytokine CXCL10. Validation of the altered ISG expression by RT-PCR (**Fig. 5E, S5C**) demonstrates that loss of hnRNPM results in significant upregulation of ISGs.

To examine whether the ISG upregulation in hnRNPM KD cells is indeed the effect of dsRNA-mediated IFN-I response, we depleted three dsRNA sensors DDX58, IFIH1, and TLR3 in hnRNPM KD cells (**Fig. S5D**) and determined whether loss of dsRNA sensors mitigates ISG induction (**Fig. 5F**). As shown in **Fig. 5G**, control cells express very low levels of ISGs. Knockdown of dsRNA sensors in control cells does not affect ISG levels. In contrast, hnRNPM KD cells express high levels of ISGs and their expression is significantly reduced when dsRNA sensors are knocked down, showing that dsRNA sensing is required for ISG upregulation in hnRNPM KD cells (**Fig. 5G**).

As a complementary approach, we cloned the regions of cryptic exons of NUDT19 and MED15 that are predicted to form dsRNAs in an expression plasmid and ectopically expressed them in 293FT cells (**Fig. 5H**). We detected a significant upregulation of ISGs in cells transfected with the NUDT19-or MED15-cryptic exon expressing plasmid, supporting our findings that hnRNPM-repressed cryptic exons form dsRNAs, which in turn stimulates IFN-I-mediated ISG upregulation (**Fig. 5I**).

### Tumors with low hnRNPM expression show upregulated IFN-I responses and better survival of cancer patients

Elevation of the IFN-I pathway is widely observed to activate anti-tumor immunity.^54–56^ Given that loss of hnRNPM stimulates IFN-I response through cryptic splicing-induced dsRNA formation, we examined whether hnRNPM expression in tumors is associated with IFN-I signatures. Using the TCGA database, we separated tumors into hnRNPM^high^ and hnRNPM^low^ groups based on hnRNPM normalized expression levels and conducted GSEA analyses across all available cancers in TCGA by focusing on 50 “hallmark” gene sets from the Molecular Signature Database (MSigDB, **Fig. S6A**). Strikingly, in as many as 16 out of 33 cancer types, hnRNPM^low^ tumors showed significant enrichment of immune pathways, especially those associated with IFN-I response, such as IFN-α, IFN-γ, and IL6/Jak/Stat3 pathways (**Fig. 6A**). These hnRNPM^low^ tumors also exhibited disadvantages in cell proliferation (**Fig. 6A**). These results from TCGA datasets are reminiscent of our findings in hnRNPM KD cells (**Fig. 5C, S5A**), and are further validated using the CPTAC3 dataset (**Fig. S6B**). Additionally, we stratified breast tumors by its four subtypes (Luminal A, Luminal B, HER2, Basal), and found that the most aggressive basal subtype has the most marked immune response upregulation in its hnRNPM^low^ tumors (**Fig. 6A**).

**Figure 6.**
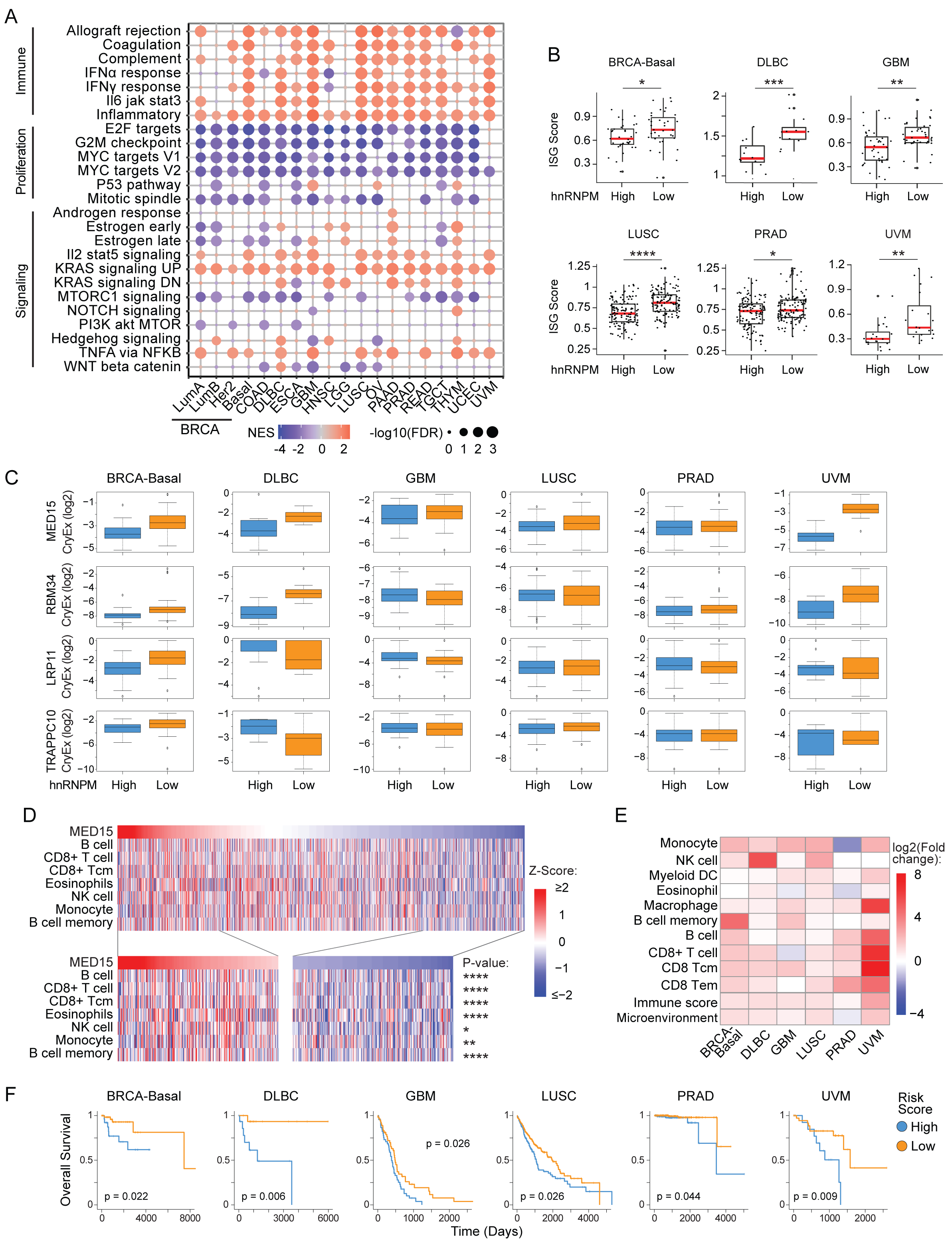
hnRNPM deficiency in cancer patients leads to upregulated IFN-I responses and better survival. **A.** Selected MsigDB hallmark gene sets enriched in hnRNPM lowly expressed tumors across TCGA cancers. The size of the circles represents the significance of enriched gene set measured by log10 transformed FDR. Color indicates normalized enrichment score for each gene set. **B.** ISG score distribution of hnRNPM lowly vs highly expressed tumors across cancers. Scores from ssGSEA analysis of ISG gene signature were computed. P values were calculated using Wilcoxon rank-sum test. **C.** Log-transformed PSI distributions of four most representative dsRNA-forming cryptic exons in TCGA. Boxplots showing an increase of PSIs in hnRNPM-low expressed tumors across cancers. **D.** A heatmap showing MED15 cryptic exon inclusion positively correlates with signatures of immune infiltration in breast cancer. Scores from ssGSEA analyses of xCell immune cell gene signatures were computed and P values were calculated between first and fourth quantiles of MED15 cryptic exon inclusion level distribution in TCGA BRCA tumors (n = 1093) using Wilcoxon rank-sum test. **E.** A heatmap showing log2 fold changes of immune cell infiltration between hnRNPM lowly and highly expressed tumors across cancers. **F.** Kaplain-Meier survival analyses showing significantly better survival for patients with low-risk scores. Risk score was calculated based on both the inclusion levels of dsRNA-forming cryptic exons in Panel 6**C** and hnRNPM expression in each tumor.

To investigate if upregulated immune responses observed in hnRNPM^low^ tumors are associated with dsRNA-stimulated viral responses, we directly compared ISG expression in hnRNPM^high^ and hnRNPM^low^ tumors. We performed single-sample GSEA (ssGSEA)^57^ to assign ISG scores to each tumor and found that six types of cancers, including the basal subtype of breast cancer, exhibited significantly increased ISG scores in hnRNPM^low^ tumors (**Fig. 6B**). These results are recapitulated using CPTAC3 datasets (**Fig. S6C**), further confirming our findings that loss of hnRNPM results in ISGs upregulation. Moreover, we evaluated four hnRNPM-repressed dsRNA-forming cryptic splicing events (MED15, RBM34, LRP11, TRAPC10) in tumors and observed an overall upregulation of cryptic exon inclusion in hnRNPM^low^ tumors (**Fig. 6C**). These results are congruent with our in vitro findings that loss of hnRNPM produces cryptic splicing-mediated dsRNAs, stimulating interferon responses.

Upregulated interferon responses facilitate recruitment of immune cell subtypes, such as NK cells and CD8^+^ T cells, which elicit anti-tumor immunity.^58–60^ Using individual immune cell gene signatures curated from xCell,^61^ we assigned immune cell infiltration scores to each tumor through ssGSEA. By ranking TCGA breast tumors by their PSI values of MED15 cryptic intron retention (**Fig. 6D**, left to right), we found that MED15 PSI values are positively correlated with immune infiltration for a set of selected immune cells, including NK cells and CD8^+^ T cells. Encouraged by our findings, we inspected immune cell infiltration in hnRNPM^high^ and hnRNPM^low^ tumors across the above six cancer types. Remarkably, many innate and adaptive immune cells known to exhibit anti-tumor functions, including NK cells and CD8^+^ T cells, are highly infiltrated in hnRNPM^low^ tumors (**Fig. 6E**), supporting the phenotype of IFN-I and ISG upregulation in hnRNPM^low^ tumors.

Lastly, we examined whether the findings presented here may be associated with the survival of cancer patients as anti-tumor immunity response improves patient survival. We built a multi-variate survival regression model by taking into consideration both hnRNPM expression and dsRNA forming cryptic exon inclusion rates. We extracted a composition score (a.k.a risk score) from the model for each tumor and classified the tumors into low-and high-risk groups. Kaplan-Meier survival estimation revealed that patients with hnRNPM cryptic splicing-based low-risk scores show better survival (**Fig. 6F**). Altogether, our results reveal that tumors with lower hnRNPM expression show higher tendency of cryptic splicing, increased levels of ISGs and IFN-response, and elevated immune cell infiltration in tumors, which ultimately results in better patient survival.

## Discussion

It has been widely observed that human introns contain an enormous amount of pseudo splice sites whose sequences are indistinguishable from canonical splicing sites. To maintain the accuracy of pre-mRNA splicing, cells must employ a powerful mechanism to repress pseudo splice sites from recognition to suppress cryptic splicing. The present study has shed light onto an RBP-mediated mechanism for cryptic splicing suppression, which is crucial to maintaining transcriptome integrity.

Through surveying RBPs that preferentially bind to deep introns, we identified hnRNPM as an essential cryptic splicing suppressor. We show that hnRNPM binds near pseudo splice sites that are enriched in LINE elements. Loss of hnRNPM triggers IFN response at least in part through cryptic splicing produced dsRNAs, a phenotype that resembles host response upon viral infection. In cancer patients, dsRNA-forming cryptic splicing in hnRNPM lowly expressed tumors is associated with elevated IFN response and tumor immune surveillance, providing a new connection between RBP loss, dsRNA production, and tumor immunity.

hnRNPM is a ubiquitously expressed nuclear RBP with typical RNA recognition motifs, and it contributes significantly to nucleic acid metabolism including splicing.^62, 63^ It is well established that hnRNPM generally serves as a repressor of alternative splicing at proximal introns.^31^ For example, hnRNPM represses a large number of alternative splicing events associated with epithelial cells.^31^ Consequently, loss of hnRNPM binding promotes EMT and cancer metastasis through increased inclusion of mesenchymal alternative exons.^31^ In this study, we found that hnRNPM is also a repressor of cryptic pseudo-exon inclusion. Upon the loss of hnRNPM, we detect over a thousand cryptic exons strongly regulated by hnRNPM. A majority of cryptic exon-containing protein-coding transcripts are potential NMD targets, due to a frameshift which may introduce a pre-mature stop codon. Additionally, cryptic exon-containing transcripts can potentially be translated into novel proteins, which may affect cell fitness and produce neo-antigens in tumors.^64–66^ These findings shine light on the role of hnRNPM not only as a repressor of alternatively spliced and cryptic exon inclusion, but also a guardian of transcriptome stability and proteome integrity.

We discovered that hnRNPM-regulated cryptic exons are highly hnRNPM-bound and are specifically enriched in deep intronic LINEs. Some of the long cryptic exons are replete with LINEs together with inverted Alu elements scattered in between, which can form long dsRNAs. These cryptic exon-derived dsRNAs initiate antiviral defense pathways by mimicking a viral infection. Likewise, in tumors with low hnRNPM expression, inclusion of dsRNA-forming cryptic exons are increased, along with an overall upregulation of interferon responses, indicating that viral defense pathways in hnRNPM-low tumors may be activated through dsRNA upregulation. Previous studies on the loss of hnRNPM in macrophages indicates that hnRNPM represses innate immune gene expression through alternative splicing of the IL-6 gene^67^ and it has a negative regulating function on the recognition of viral RNA by interacting with RIG-I-like receptors (RLRs).^68^ Our findings show that repeat elements inside hnRNPM-repressed cryptic exons can form dsRNAs and that these dsRNAs can stimulating cellular interferon responses. This mechanism provides a novel source for dsRNA-induced immune response upregulation.

Based on our data, not all cryptic exons are suppressed by one single RBP. Through analyses of a variety of RBPs regulating cryptic splicing using ENCODE eCLIP and RNA-seq data, we discovered that MATR3, SUGP2 and hnRNPM share over a quarter of their repressed cryptic exons. Knocking down MATR3 or hnRNPM alone can have the same effect on some of their overlapped cryptic exons (Data not shown). With the cryptic exon identification pipeline we developed, future work on how RBPs can combinatorically repress cryptic splicing would help paint a more comprehensive portrait of cryptic splicing biology.

Our findings on cryptic exons in cell lines and tumors can be applied to the study of normal tissues and development as well, since cancer can be perceived as abnormal development in human tissues. We believe that completing a thorough exploration of the cryptic exon landscape changes in cells under various normal conditions would be as critical of a breakthrough for basic research in biology as our previous decades of dedication on understanding alternative splicing biology. Cryptic exons can be a “suicidal activation code” embedded in genomes by evolution that is triggered only when cells are in danger (e.g., fluctuation of essential RBPs in cells) and its unanticipated function in stimulating cellular innate immunity is such a sophisticated design by nature that could potentially revolutionize our future cancer treatment. Furthermore, our discovery of the primate-specific property of hnRNPM-regulated cryptic exons has intrigued us to study cryptic exons in various species other than human. Notably, our established cryptic exon identification pipeline is not limited to the human genome only. This would give us more confidence in unveiling long-overlooked cryptic exon perspectives about evolution and differential disease predisposition in various animals due to their genomic composition discrepancies that may be heightened in cryptic splicing.

There has been a plethora of treatments based on targeting various proteins in cells to trigger anti-tumor immunity over the years.^69, 70^ Here, we discovered an unprecedented mechanism in which depleting hnRNPM can elevate antiviral signaling in tumors via cryptic exon-derived long stem-like dsRNAs, triggering a type I interferon response. Such immunity-enhancing properties of hnRNPM-regulated cryptic exons possess vast clinical potential and may expand horizons of treatment for diseases other than cancer. Recently, RBPs are emerging as therapeutic targets for prevention and treatment of diseases related to aberrant alterations in genomes such as cancer.^71–73^ hnRNPM inhibition or targeting dsRNA-forming cryptic exon splicing may be intriguing new therapeutic methods for activating immunity in patients.

## METHOD DETAILS

### Cell lines and cell culture

Human embryonic kidney cell line 293FT, cervical cancer cell line HeLa, breast carcinoma cell line-derived LM2 and BM2 cells were grown in DMEM supplemented with 10% FBS, L-glutamine and Pen/Strep at 37 ^0^C. HMLE cells were grown in Mammary Epithelial Cell Growth Medium (Lonza, USA). All cells were passaged every 2 – 3 days.

### iCLIP assay and analysis

Two biological replicates of HMLE cells were UV crosslinked and iCLIP was performed as previously described using anti-hnRNPM antibody (Origene).

Data analysis was conducted using the Clip Tool Kit (CTK) v1.0.3. After processing and mapping iCLIP reads in CTK, replicates were pooled and single nucleotide resolution binding sites for hnRNPM were determined using the following strategy: 1) Statistically significant iCLIP peaks were called and delimited by one half the peak height (adjusted p < 0.05); 2) Significant crosslinking induced truncations sites (CITS) were called (adjusted p < 0.05) using the CTK CITS.pl script; 3) CITS were further filtered by retaining only those that overlapped with a significant peak; 4) CITS were extended upstream and downstream by 10 nucleotides to identify local binding sites for further analysis.

De novo motif analysis was performed using HOMER v4.10 findMotifsGenome.pl script with the following non-default parameters -p 4 -rna -S 10 -len 4,5,6 -size 100 -chopify. De novo motifs were computed compared to a background of shuffled human introns.

For the GU-rich hnRNPM binding motif RNA map analysis, 14 GU-rich 5-mer motifs used in a previous study to identify hnRNPM binding motifs were used to screen for motif enrichment.^74^ The motifs were = [‘UGUGU’, ‘GUGUG’, ‘UUGUG’, ‘GUGUU’, ‘UGUUG’, ‘UGUGG’, ‘GUUGU’, ‘GGUGU’, ‘UGGUU’, ‘UUGGU’, ‘UGGUG’, ‘GUGGU’, ‘GUUGG’, ‘GGUUG’].

### Calculation of a normalized binding score for RBPs on different gene features

To compare binding preferences of RBPs across different gene features, a normalized binding score for each RBP on each gene feature was calculated as follows. RBPs with more than 5% of their reproducible and significantly enriched peaks located in introns in both HepG2 and K562 cells were selected for further analysis.^27^ Number of each RBP’s binding sites at each gene feature was normalized against each RBP’s overall number of binding sites across pre-mRNA, generating a binding metric at different gene features in each RBP. The metric was then normalized by the genomic length of its associated gene feature (5’UTR, exon, 3’UTR, proximal (PROX) intron, and deep intron). Enrichment on a specific feature is defined by normalizing to the mean between the features.

### Calculation of a preferential binding metric for hnRNPM across pre-mRNA

Significant hnRNPM iCLIP binding peaks in HMLE cells were calculated for the percentage of peaks counts at each gene feature and divided by the corresponding genomic length of each gene feature. Number of significant hnRNPM iCLIP binding peaks of hnRNPM in the rest of gene features was divided by the total genomic length of the rest gene features. hnRNPM preferential binding metrics were calculated by dividing the two divisions above (each gene feature/the rest gene features), then log2-transformed and visualized across pre-mRNA.

### Deep RNA sequencing and data analysis

Three biological replicates for control and hnRNPM knockdown HMLE cells were collected with 1ml TRIzol for a 10cm dish. RNA samples were extracted following by the TRIzol Reagent kit from Invitrogen. The purified RNA samples were submitted to the Genomic Facility at University of Chicago for RNA quality validation, RNA-seq library generation and paired-end sequencing on HiSeq 4000. RNA-seq reads were aligned to the human genome (GRCh37, primary assembly) and transcriptome (Gencode version 24 backmap 37 comprehensive gene annotation) using STAR v2.7.9a ^75^ with the following non-standard parameters --outSAMstrandField intronMotif -- outFilterType BySJout --outFilterMultimapNmax 1 --alignSJoverhangMin 8 -- alignSJDBoverhangMin 3 --alignEndsType EndToEnd. Only uniquely aligned reads were retained for downstream analysis.

Differential gene expression analysis was performed by counting reads over genes from the same annotation as alignment using featureCounts version 1.5.0 with the following non-default parameters -s 0 -a. Differential gene expression analysis was conducted using DESeq2^76^ performed on genes with at least 5 counts present in at least half of the samples. Significantly regulated genes were defined as genes with an | log2FC | > 2 and FDR < 0.05.

Differential alternative splicing was quantified using rMATS version 4.0.2 with the following non-default parameters –readLength 100 –cstat 0.01 –libType fr-secondstrand. To identify significant differential splicing events, we set up the following cutoffs: FDR<0.05, |ΔPSI|>=0.1, and average junction reads per event per replicate >=20.

Gene set enrichment analysis (GSEA) was conducted through our in-house script that uses fgsea^77^ on genes ranked by log2FC with 1000 gene set level permutations and considers only gene sets of size 15 to 500 genes. Significantly enriched pathways were selected by FDR < 0.05. Gene ontology (GO) biological process analysis was conducted using http://webgestalt.org with default setting^78^. By ranking log2FC of hnRNPM knockdown vs control, the top 300 upregulated and downregulated genes were uploaded for the analysis separately.

### De novo identification of hnRNPM-regulated cryptic exons and retained introns

To predict exons from RNA-seq data, STAR v2.7.9a was used to generate BAM files that are sorted by coordinates and used Stringtie v2.1.5 to annotate any expressed exons for each sample. All exons that were not identical with exons annotated in GENCODE v19 were referred to as ‘cryptic’. To narrow down to adequately expressed cryptic exons that arise from introns, ‘cryptic’ exons which are annotated exons with read alignments extending across the 5’ss, 3’ss, or both ends and have at least 5 reads aligned to their spliced junctions were kept. Full-length intron retention events are analyzed separately. hnRNPM-regulated cryptic exons are defined as events with a PSI value less than 0.1 in control cells and ΔPSI values (ΔPSI = PSI{KD} – PSI{Ctrl}) larger than 0.1. hnRNPM-regulated retained introns, PSI value in control cells should be less than 0.1 and the PSI ratio (PSI ratio = PSI{KD} / PSI{Ctrl}) larger than 2.

### RT-PCR analysis

RT-PCR and semi qRT-PCR were conducted using RNA extracted from cells processed with the E.Z.N.A. Total RNA Kit (Omega Bio-Tek). RNA concentration was measured using a Nanodrop 2000 (Thermo Fisher Scientific). cDNA was generated via reverse transcription using the GoScript Reverse Transcription System (Promega) with 1 ul GoScript RT and 250 ng of RNA in a total volume of 20ul followed by incubation at 25 °C for 5min, 42 °C for 30min, and 70 °C for 15min.

qRT-PCR was performed using GoTaq qPCR Master Mix (Promega) per the manufacturer’s instructions on a CFX Connect Real-Time PCR system (BioRad) using a two-step protocol and supplied software. For every qPCR sample per biological replicate, two technical replicates were performed, and Ct counts were averaged. Normalization and quantification of qPCR data was done using the 2^−ΔCt^ method relative to TATA-binding protein (TBP) expression (Livak and Schmittgen 2001). Primer specificity was verified with melt-curve analysis during qPCR.

Primers for semi-qRT-PCR analysis were designed on constitutive exons flanking each variable exon.^79^ Semi-qRT-PCR generates both exon inclusion and skipping products in one PCR reaction, which were separated through agarose gel electrophoresis. PCR product intensity was quantified on a QIAxcel Advanced System. Primers for qPCR and semi-qRT-PCR are included in Supplemental Table S3.

### Western blot

Cell lysates from HMLE shLuc and shM2 were separated by 4-20% SDS-PAGE (Genscript) and transferred to a PVDF membrane (10600023 GE). Membranes were blocked with 5% milk in 1x TBST for 30 minutes and then incubated in primary antibodies overnight at 4°C. Primary antibodies used were hnRNPM (Origene technologies, TA301557, 1:50,000), GAPDH (EMD Millipore, MAB374, 1:3,000), and β-actin (Sigma, A5441, 1:3,000). Membranes were then washed in TBST and incubated in the corresponding HRP-conjugated secondary antibody for 1 hour at room temperature. After final TBST washing, membranes were imaged using a ChemiDoc (BioRad) with Immobilon Western Chemiluminescent HRP Substrate (Millipore).

### Splicing minigene reporter and assay

ADIPOR2 and MED15 cryptic exons and 500 bp of their up-and downstream intronic regions were amplified from genomic DNA of HMLE cells using NEB Phusion High-Fidelity DNA Polymerase (M0530). Cloning primers for amplifying ADIPOR2 cryptic exon and proximal introns are: Forward: 5’-TAAGCAGGATCCTGCGTATGTTGAACCAGCCTTG-3’; Reverse: 5’- TAAGCA GGATCCTGAATCCAGGAGCTGGTTTTCTG -3’. Cloning primers for amplifying MED15 cryptic exon and proximal introns are: Forward: 5’- TAAGCAGGATCCCACAGCTACTGTACTGAA GCCTTG-3’; Reverse: 5’- TAAGCAGGATCCTCCTCTCAACAAGCACAAGCTG-3’. PCR products were purified with IBI Scientific Gel/PCR DNA Fragment Extraction Kit (IB47020). The pETv8 backbone and PCR products were digested with BamHI restriction enzyme (NEB) and gel purified. The cut backbone and PCR products were ligated together using NEB T4 DNA Ligase (M0202). Plasmid sequences were confirmed by DNA sequencing.

2.5×10⌃5 293FT cells were seeded in each well of 24-well plate 20-24 hours prior to transfection.100 ng splicing minigene and increasing amount of pcDNA3-control or pcDNA3-hnRNPM plasmid^31^ were cotransfected into 293FT cells using Lipofectamine 2000 (Invitrogen). Cells were collected 24 hours after transfection for RNA extraction and analysis.

### Cytoplasmic and nuclear fractionation for RNA isolation

293FT cells growing in a 15cm tissue culture dish were collected in PBS using a tissue culture scraper. After two washes with PBS, cells were centrifuged and resuspended in 1X hypotonic buffer (20 mM Tris-HCL (pH 7.4), 10 mM NaCl, 3 mM MgCl2). After 15 minutes incubation on ice, 10% NP-40 was added, and the vial was vortexed for 10 seconds. The fractions were separated by centrifugation for 10 minutes at 3,000 rpm at 4°C. The supernatant containing the cytoplasmic fraction was collected, and the RNA was extracted using TRIzol LS Reagent (Invitrogen). The pellet containing the nuclear fraction was used for RNA pull-down.

### dsRNA overexpression

The dsRNA regions for NUDT19 and MED15 were amplified from cDNA of hnRNPM KD HMLE cells using NEB Phusion High-Fidelity DNA Polymerase (M0530). Primers used to amplify the NUDT19 dsRNA-coding region are: Forward: 5’- AGAGGTCTGTTACCACTCTTGCAGG-3’; Reverse: 5’- ACTGCACCCAGCCAAGTTCCAA-3’. Primers used to amplify the MED15 dsRNA-coding region are: Forward: 5’- CCTATAAAACCAAGACATGCAGGCAC-3’; Reverse: 5’- GGACTGTTTGAAAATGAGATGC ATGGC-3’. PCR products were run through an agarose gel and purified with IBI Scientific Gel/PCR DNA Fragment Extraction Kit (IB47020). A second PCR was performed using the PCR products and restriction enzyme flanking primers: NUDT19 cloning Forward: 5’- CGCTCTAGCCCGGGCGGATCCAGAGGTCTGTTACCACTCTTGCAGG-3’; Reverse: 5’- GTCATCCTTGTAGTCCTCGAGACTGCACCCAGCCAAGTTCCAA-3’. MED15 cloning Forward: 5’- CGCTCTAGCCCGGGCGGATCCCCTATAAAACCAAGACATG CAGGCAC- 3’; Reverse: 5’- GTCATCCTTGTAGTCCTCGAGGGACTGTTTGAAAATGAGATGCATGGC-3’. The pBRIT backbone and PCR products were digested with NEB restriction enzymes BamHI and XhoI for 3 hours and then gel purified. The backbone and PCR inserts were ligated together using NEB T4 ligase (M0202) according to the product protocol with a 1:6 molar ratio. Plasmid sequences were confirmed by DNA sequencing.

293FT cells were seeded into a 24-well plate at 5×10⌃4 cells per well. The following day, 500 ng plasmid was transfected using Lipofectamine 2000 (Invitrogen) according to the manufacture instruction. Media was replaced 24 hours after transfection. RNA was collected 48 hours after transfection and purified using E.Z.N.A. Total RNA Kit I (R6834 Omega BioTek).

### siRNA Knockdown

Cells were plated on a 24-well tissue culture treated plate at 5-7×10⌃4 cells per well. The following day, 20-60 nM siRNA was transfected using Lipofectamine RNAiMAX reagent (Invitrogen). Media was replaced after 24 hours. RNA was collected 48 hours post-transfection using E.Z.N.A. Total RNA Kit I (Omega BioTek). siRNA sequences are provided in Supplemental Table S3.

### RNA pull-down

RNA oligonucleotides labeled with biotin at the 5’-end were synthesized by Integrated DNA Technologies. The RNA sequences used in this study are listed as follows: ADIPOR2 Oligo 1: UUUCUGUGGGAUUGGUGGUA, ADIPOR2 Oligo 2: UACUUUGUAUUUCUGUGGGA, TRAPPC10 Oligo: CUUCUGCUUGUUUGUGACCC. 400 pmol Biotinylated RNA oligos were conjugated with 50 μl of streptavidin beads (50% slurry; ThermoFisher) in a total volume of 300 μl of RNA-binding buffer (20 mM Tris, 200 mM NaCl, 6 mM EDTA, 5 mM sodium fluoride and 5 mM β-glycerophosphate, PH 7.5) at 4 °C on a rotating shaker for 2 hours. After washing three times with RNA-binding buffer, RNA-beads conjugates were incubated with 100 μg of nuclear extracts in 500 μl RNA-binding buffer at 4 °C on a rotating shaker overnight. Beads were then washed with RNA-binding buffer for three times and the RNA pull-down samples were eluted with 2 × SDS loading buffer for western blot analysis.

### Identification and analysis of RNA editing sites

Identification of RNA editing sites from RNA-seq reads was performed according to our previously published work.^80^ In brief, RNA-DNA differences (RDDs) were identified from RNA-seq reads.^81^ The identified RDDs were required to be covered by 5 or more reads, at least 2 edited reads, and 10% editing ratio. RDDs should also not be located in homopolymers, splice sites, dbSNP database, and simple repeats.^82^ The GIREMI tool was used to further select editing sites based on their mutual information with genetic variants.^83^ To identify differentially edited sites between the two conditions, we used the parametric test for allelic bias as previously described.^84^ Site-specific normal distributions were parameterized by the mean editing level between replicates and the expected variance calculated from the mean coverage. The FDR was calculated using the Benjamini-Hochberg method. Differentially edited sites were called by requiring FDR ≤ 10% and the absolute change in the editing level between knockdown and control ≥5

### RNA fluorescent in situ hybridization for cryptic exons

The Stellaris RNA FISH probe designer was used to generate 45-48 probes labelling constitutive exons and cryptic exons of TRAPPC10 and MED15 (Supplemental Table S4). For visualization, exon probes were coupled to Quasar670 and cryptic exon probes to Quasar570. HMLE control and hnRNPM KD cells were fixed in PBS/4% PFA for 30 min at room temperature and RNA staining was performed according to the stellaris RNA FISH protocol for adherent cells. Images were acquired using a GE Healthcare DeltaVision LIVE High Resolution Deconvolution Microscope and analyzed in Fiji/ImageJ.

### Metaplot and motif enrichment

For the CLIP binding metagenes, the 2 kB up-and downstream intronic sequences flanking the cryptic exons were also obtained. For each of the three following types of exons: hnRNPM-regulated cryptic exons (CryEx), alternatively spliced exons, and annotated background exons, we padded their coordinates with 2 kB up-and downstream. We then intersected them with the hnRNPM iCLIP binding sites using bedtools. The “% exon” represents the number of exons containing an overlapping binding site at a given position over the total number of exons. The equation was as follows:

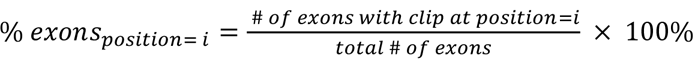

The binding probability (% exons) was further smoothed by using a moving average of 500-bases.

The motif score is defined as the mean percentage of bases containing any of the hnRNPM motifs in a 50-bases window. The score for a given position is the percentage calculated for the window ending at that position. The equation as follows:

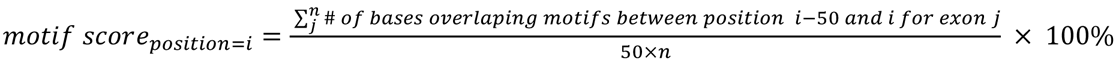

### RNA structure prediction

RNA structure prediction was done using the RNAfold web server (http://rna.tbi.univie.ac.at/cgi-bin/RNAWebSuite/RNAfold.cgi) with default options.

### ISG score calculation

ISG gene signature was curated using MsigDB hallmark interferon alpha and beta gene sets (https://www.gsea-msigdb.org/gsea/msigdb/collections.jsp) and ISGs induced by dsRNAs in HeLa cells 6 or 12 h after transfection. ssGSEA was conducted using ISG gene signature and its calculated score is represented as ISG score. The higher the ISG score the higher the interferon responses in cells.

### Immune infiltration score calculation

The infiltration level of different immune cell populations were estimated by ssGSEA^57^ in the R Bioconductor package Gene Set Variation Analysis (GSVA, v1.44.3) using default parameters. Gene sets from published xCell paper^61^ were curated and used to derive ssGSEA scores. Calculated ssGSEA scores are used to represent immune infiltration scores.

### Clinical and molecular data of TCGA study

Gene expression data was downloaded from the GDC portal (https://portal.gdc.cancer.gov). After filtering out low abundance genes that have less than 5 reads across half of the samples in all TCGA cancer types, we applied log2-transformed counts per million (CPM) normalization and the trimmed mean of M-values (TMM) adjustment on count matrix for each cancer. We then defined 1^st^ quartile of hnRNPM expressed tumors as the hnRNPM-low tumors, and 4^th^ quartile of hnRNPM expressed tumors as the hnRNPM-high tumors.

### Multivariate survival model and risk score calculation

Four representative dsRNAs containing cryptic exons from MED15, RBM34, TRAPPC10, and LRP11 genes were chosen for multivariate survival analyses. By sliced binary alignment map (BAM) files of cryptic splicing regions in these genes from TCGA, we calculated PSI values for these cryptic exons for each tumor across cancer types: BRCA-Basal, DLBC, GBM, LUSC, PRAD, and UVM. We then fit a multiple Cox regression model with the above-mentioned four cryptic exon PSIs and hnRNPM expression levels. Risk scores were extracted from the model and assigned to each tumor. Subsequently, all samples from each cancer were divided into high-or low-risk groups using the cutoff of risk score means.

### Statistical analyses

All data were presented as mean ± standard deviation, unless specifically indicated. Correlation was assessed using Pearson correlation. Statistical significance tests included Fisher’s exact tests and hypergeometric tests. p-value < 0.05 was considered statistically significant. p < 0.05^85^, p < 0.01 (**), p < 0.001 (***), p< 0.0001 (****) where indicated.

### Supplemental information

Supplemental Information includes six figures and four tables.

## Acknowledgements

We thank the Cheng lab for helpful feedback on the project. We thank Dr. Wei Hong and Dr. Chao Cheng for discussions. This research was supported in part by grants from NIH R35-CA209919 (to H.Y.C), R01CA262686 and R01AG078950 (to X.X.), and R35GM131876 (to C.C). C.C. is a Cancer Prevention Research Institute of Texas Scholar in Cancer Research (RR160009).

## Author contributions

R.Z., C.C. and X.X. conceived of the project. R.Z. performed most of the bioinformatics analyses with help from C.F. and M.C. on ADAR analysis, S.H. on CLIP analysis, G.V. on CLIP metaplot analysis and T.C. for TCGA bioinformatics supervision. M.D., J.L., G.B., and A.V. performed the experiments. R.F. and H.C. performed CLIP. C.C. and X.X supervised the study. R.Z. and C.C. wrote the manuscript and X.X. helped with the editing. All contributors read the manuscript and provided extensive inputs.

## List of Supplementary files and Tables

Figure S1. Related to Figure 1: hnRNPM iCLIP on HMLE cells.

Figure S2. Related to Figure 2: Quantitative characterization of splicing changes upon hnRNPM knockdown.

Figure S3. Related to Figure 3: hnRNPM binds to cryptic exons in deep introns.

Figure S4. Related to Figure 4: hnRNPM-repressed cryptic exons form dsRNAs in the cytoplasm.

Figure S5. Related to Figure 5: ISGs are upregulated upon hnRNPM knockdown via dsRNA sensing.

Figure S6. Related to Figure 6: Interferon-related responses are elevated in hnRNPM lowly expressed tumors.

Suppl. Table 1: RBP binding distribution in introns, deep introns and pre-mRNAs for ENCODE HepG2 and K562

Suppl. Table 2: PSI of top 50 representative cryptic exons across cell lines

Supple Table 3: Primers for cryptic exons, ISGs, minigene splicing events, DNA sequencing and siRNAs

Supple Table 4: Probes for RNA FISH

**Figure S1:**
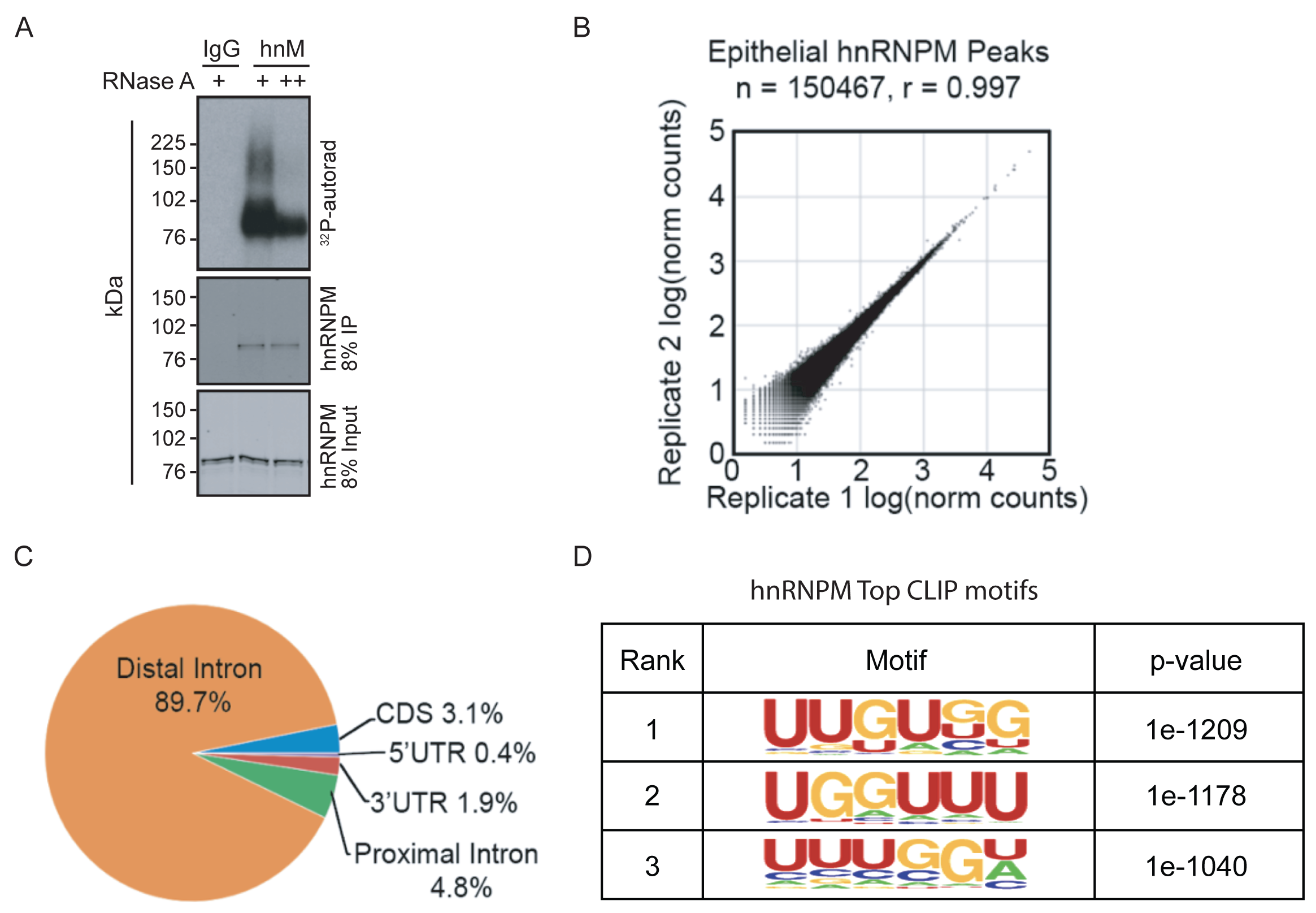
hnRNPM iCLIP on HMLE cells. **A**. An autoradiogram after crosslinking and immunoprecipitation (IP) with hnRNPM showing hnRNPM-RNA complexes, which were abolished with excess RNAse A treatment in HMLE cells. **B**. A plot showing strong correlation between hnRNPM iCLIP replicates. **C**. Genomic distribution of hnRNPM iCLIP binding sites. **D**. De novo motif analysis of hnRNPM iCLIP binding sites.

**Figure S2:**
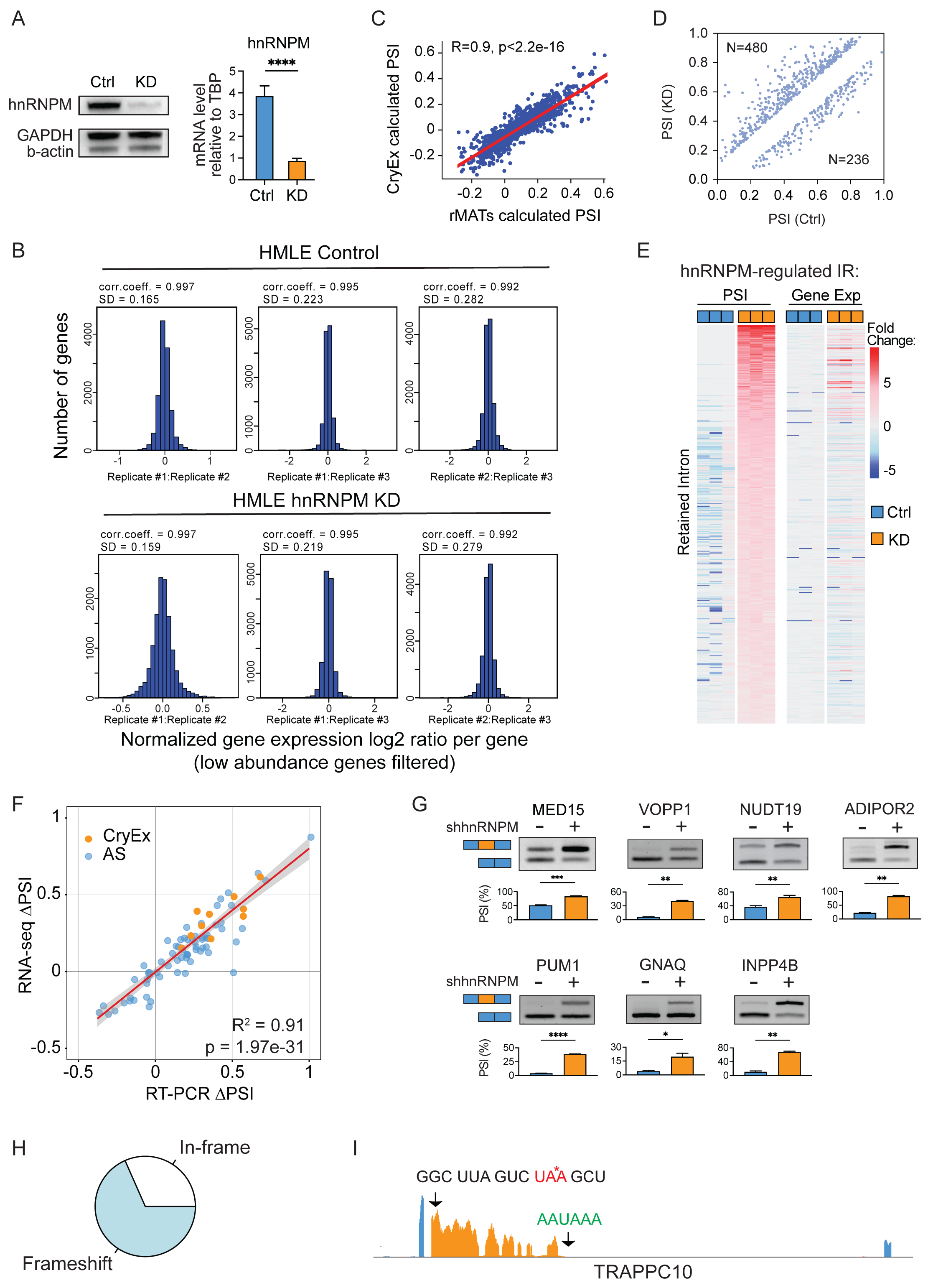
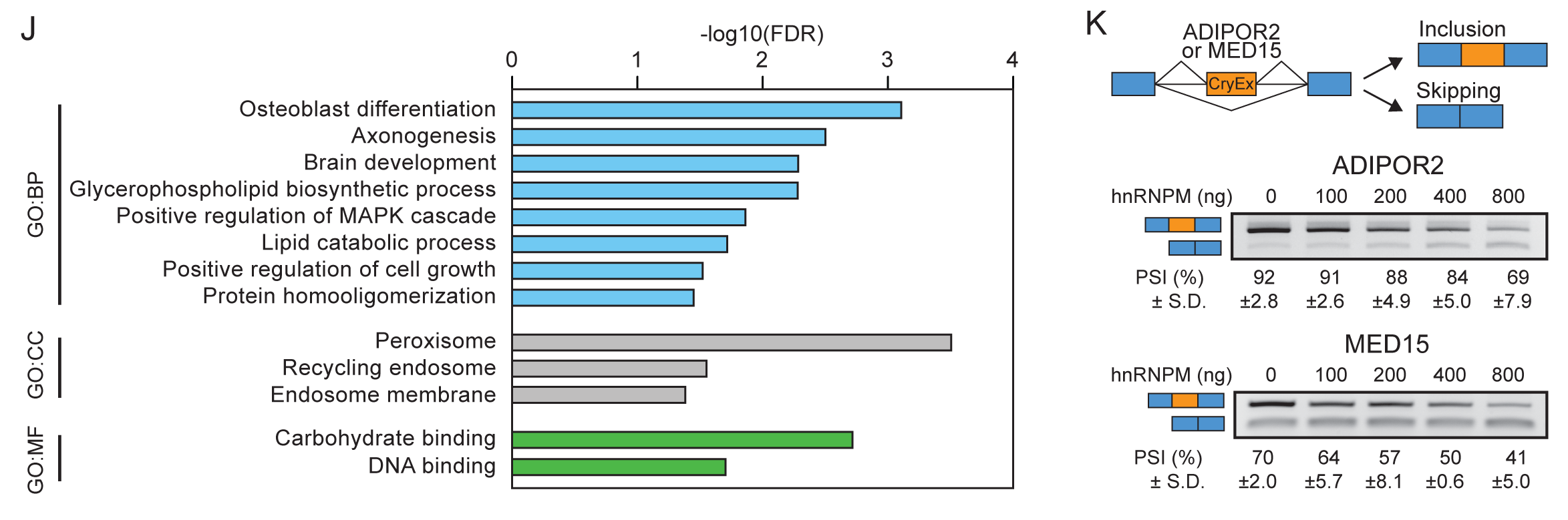
Quantitative characterization of splicing changes upon hnRNPM knockdown. **A**. Western blot and qRT-PCR showing hnRNPM knockdown efficiency. **B**. Correlation coefficients of normalized gene expression log2 ratios between replicates in HMLE control or hnRNPM KD RNA-seq samples. standard deviations (SDs) were calculated. **C**. Strong correlation between PSI values of alternatively spliced events calculated via rMATs and our pipeline. **D**. PSI distribution of alternatively spliced events from our pipeline in hnRNPM KD vs control cells. **E.** Heatmaps showing PSI fold changes of retained introns repressed by hnRNPM and expression changes of retained intron-containing genes. **F.** High correlation of ΔPSI in selected spliced events calculated from RNA-seq and tested by qRT-PCR. Lightblue indicates alternatively spliced events and orange indicates cryptic splicing events. **G.** Semi-qPCR validation of select cryptic splicing events in **F**. **H.** Pie chart showing 73.93% cryptic exons are not multiple of three. **I.** IGV plot showing potential STOP codon and polyA site in the cryptic exon of TRAPPC10. **J.** GO analysis on genes that contain hnRNPM-repressed cryptic exons. **K.** A diagram showing the design of minigene plasmid introduced with hnRNPM-repressed cryptic exons (Upper) and co-transfection experiments using minigene constructs and varying amounts of hnRNPM (Lower).

**Figure S3:**
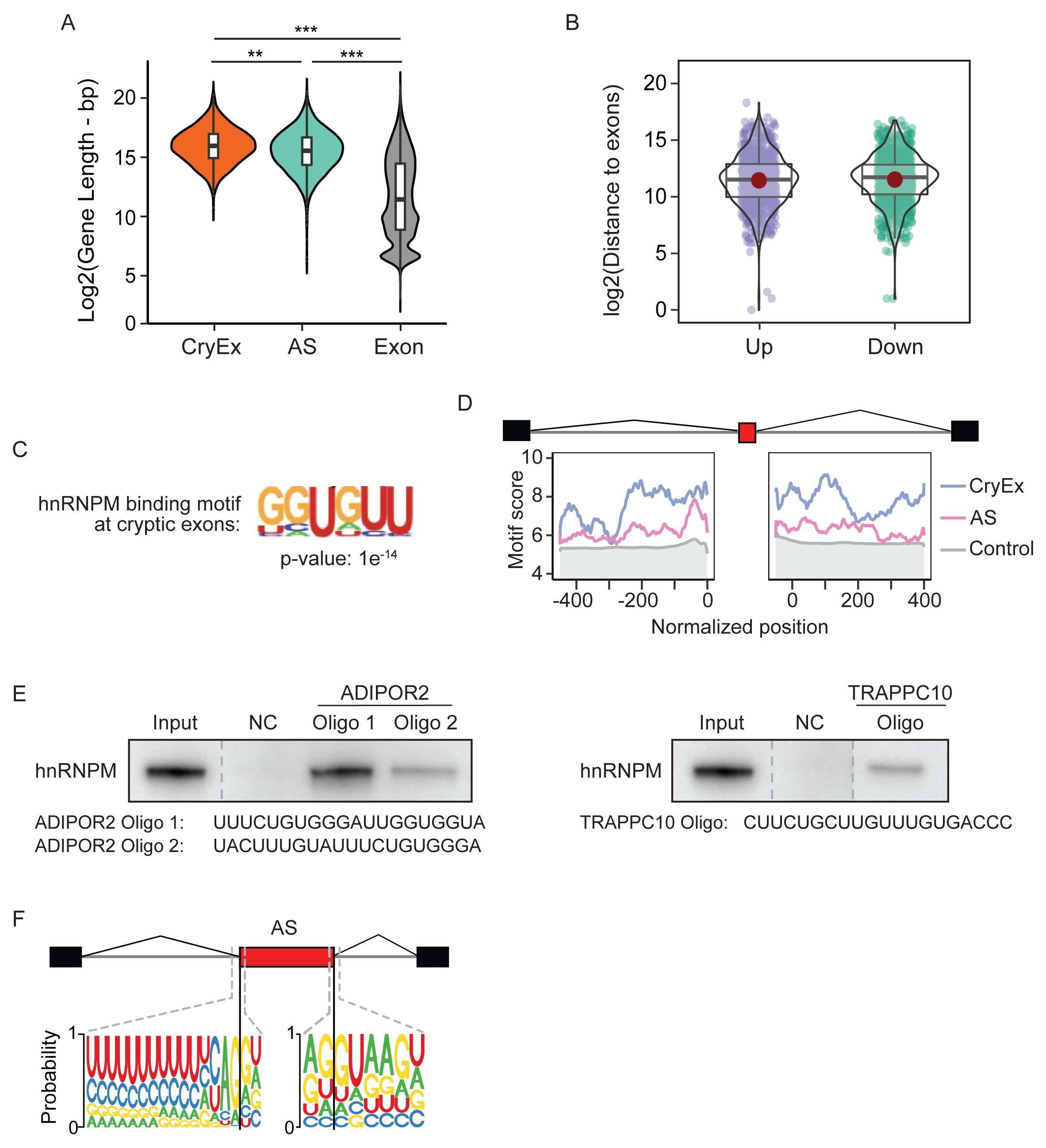
hnRNPM binds to cryptic exons in deep introns. **A**. A violin plot showing length distribution of genes containing hnRNPM-regulated cryptic exons (CryEx), alternatively spliced exons, and annotated background exons. **B**. A violin plot showing the distributions of log2 transformed distance between 3’ and 5’ splice sites of cryptic exons and their nearest upstream and downstream (Down) exons, respectively. **C**. hnRNPM binding motif enriched at hnRNPM-repressed cryptic exons. **D.** The metadata profile showing hnRNPM motif score distribution in a +/ - 2 kB window flanking three groups of exons (Upper). Metadata profile was smoothed using 50-nt bins. hnRNPM binding motif enriched at hnRNPM-regulated cryptic exons is GU-rich (Lower). **E**. WebLogo representation of consensus motifs near splice sites of hnRNPM-regulated AS exons. **F.** RNA pull-down assays showing that hnRNPM binds to cryptic exons of ADIPOR2 and TRAPPC10 genes. The sequences of cryptic exon oligos are shown at the bottom.

**Figure S4:**
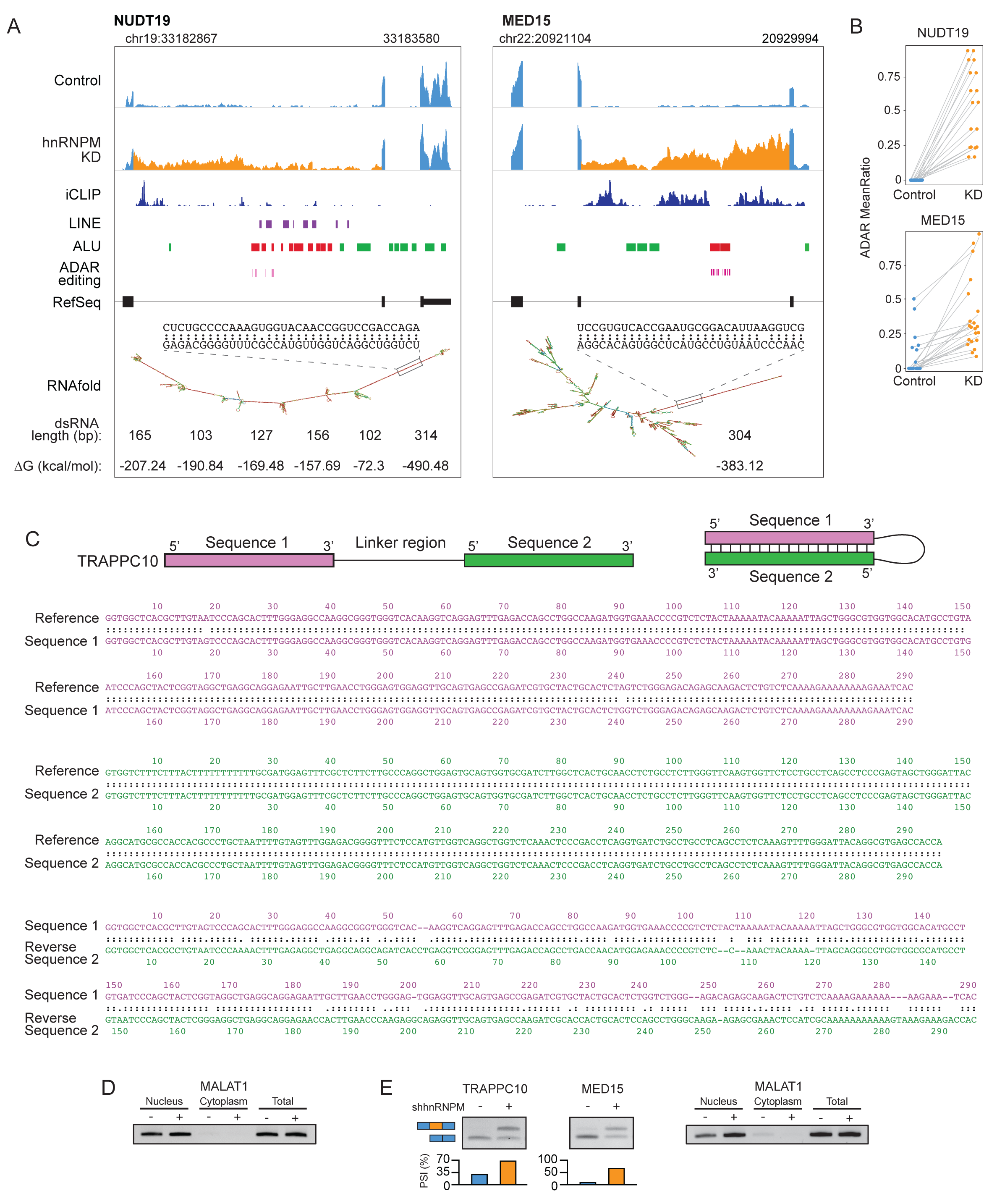
hnRNPM-repressed cryptic exons form dsRNAs in the cytoplasm. **A**. Representative examples of dsRNA-forming cryptic exons. Left, NUDT19. Right, MED15. Top two IGV tracks indicate number of reads aligned. Third track represents hnRNPM iCLIP binding data. Purple boxes under each iCLP track are LINE elements. Green boxes are Alu elements. Red boxes are Alu elements that are predicted to form dsRNAs via RNAfold. Pink boxes are ADAR editing sites. Seventh track indicates RefSeq annotation. Eighth track represents secondary structure of cryptic exon predicted by RNAfold. Approximate lengths of highly base-paired dsRNA regions are indicated beneath the structures together with predicted folding free energies (ΔG). A fragment of the dsRNA region is indicated above the secondary structure. **B.** Dot plots showing ADAR editing in predicted dsRNA-forming regions. **C.** DNA sequencing of a predicted dsRNA-forming region in TRAPPC10 cryptic exon. Depicted on the top left are two PCR regions amplified using cDNA produced from mRNA of hnRNPM KD cells. PCR products of these regions, denoted as Sequence 1 and Sequence 2, are sequenced. Depicted on the top right is a diagram of dsRNA formation by the complimentary base-pairing. Depicted on the bottom of the diagrams are sequence alignments between ref-seq and sequenced DNAs from Sequence 1 and Sequence 2, which are nearly identical. Additionally, alignment of results from Sequence 1 and reverse sequence 2 showed they are highly matched, supporting base-pairing to form dsRNA. **D.** Semi-qRT-PCR of nuclear marker MALAT1 using cytoplasm mRNA showing no nucleus contamination. **E.** *Left:* Semi-qRT-PCR of cryptic splicing events using LM2 cytoplasmic mRNA. *Right:* semi-qRT-PCR of nuclear marker MALAT1 using LM2 cytoplasm mRNA showing no nucleus contamination.

**Figure S5:**
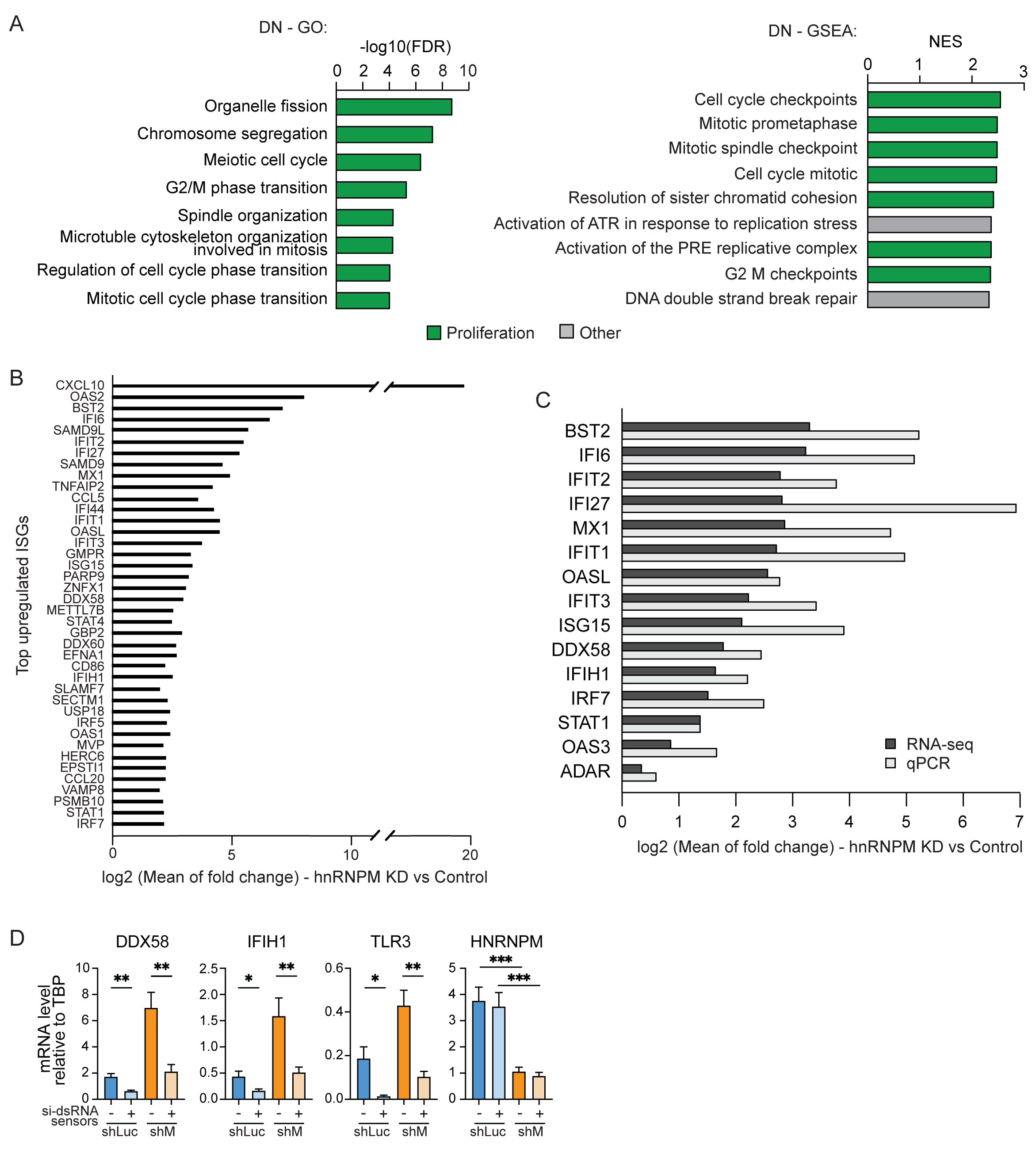
ISGs are upregulated upon hnRNPM knockdown via dsRNA sensing. **A**. Gene Ontology (Left) and Gene Set Enrichment Analysis (Right) showing top negatively enriched biological processes and pathways in hnRNPM KD cells. Cell cycle related GO terms and pathways are shown in green. Others are shown in grey. **B**. Top 40 upregulated ISGs upon hnRNPM knockdown are ranked from high to low based on their log2 transformed fold changes between mean normalized gene expression in hnRNPM KD vs control cells. **C.** Log2 transformed gene expression fold changes of ISGs from RNA-seq data and semi-qRT-PCR validation. **D**. qRT-PCR analyses showing siRNA knockdown efficiency of dsRNA sensors DDX58, IFIH1, TLR3. Compare bar 1 and 2, Bar 3 and 4. hnRNPM KD efficiency was shown on the last plot to the right.

**Figure S6:**
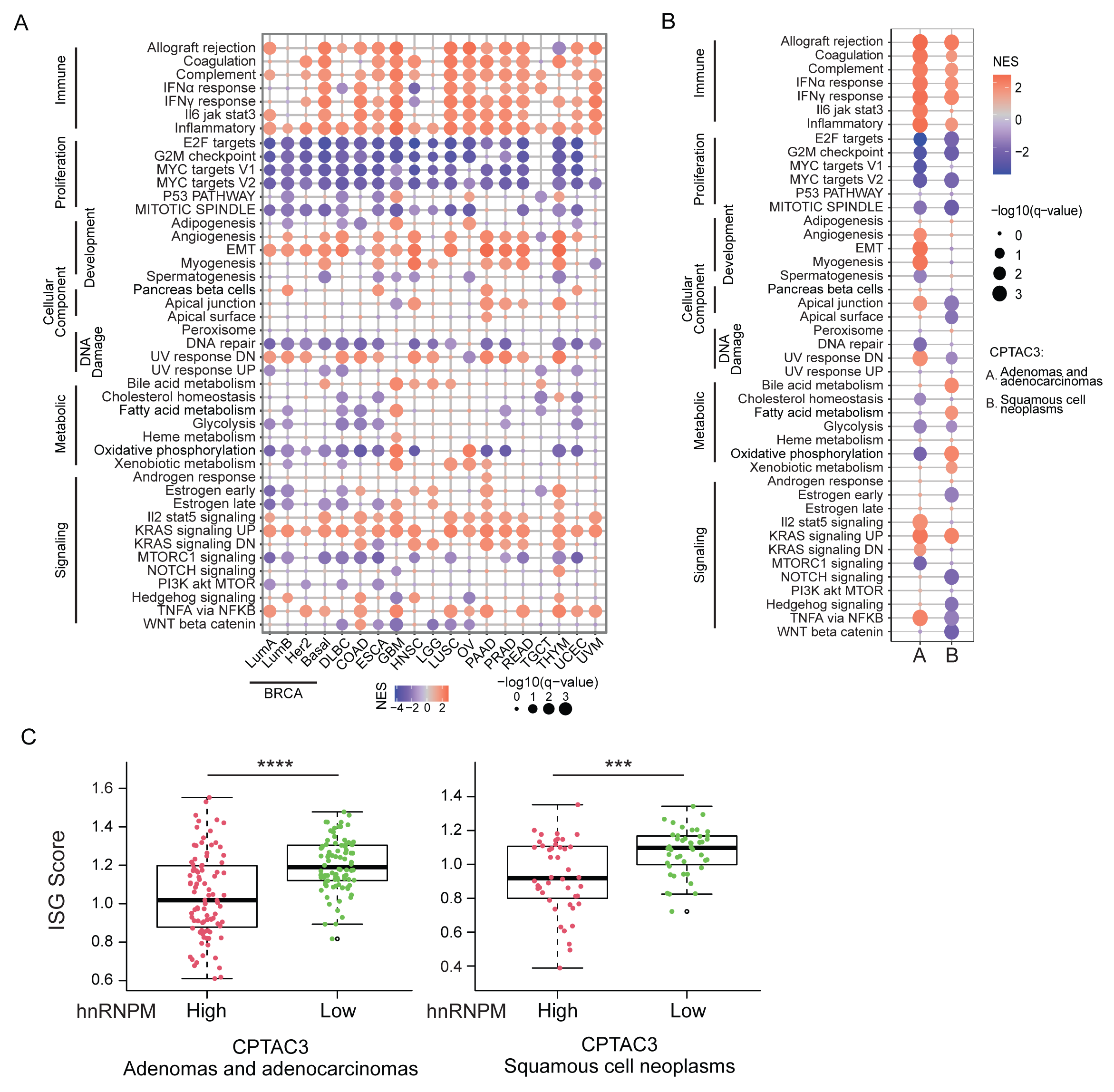
Interferon-related responses are elevated in hnRNPM lowly expressed tumors. **A.** MsigDB hallmark gene sets enriched in hnRNPM-low tumors across TCGA cancers. The size of the circles represents the significance of enriched gene sets measured by log10 transformed FDR. Color indicates normalized enrichment score for each gene set. **B.** MsigDB hallmark gene sets enriched in hnRNPM lowly expressed tumors across CPTAC3 cancers. **C.** ISG score distribution of hnRNPM-low vs hnRNPM-high expressed tumors across CPTAC3 cancers.

## References

1. Jo, B.S., and Choi, S.S. (2015). Introns: The Functional Benefits of Introns in Genomes. Genomics Inform 13, 112–118. 10.5808/GI.2015.13.4.112.

2. McCoy, M.J., and Fire, A.Z. (2020). Intron and gene size expansion during nervous system evolution. BMC Genomics 21, 360. 10.1186/s12864-020-6760-4.

3. Hare, M.P., and Palumbi, S.R. (2003). High intron sequence conservation across three mammalian orders suggests functional constraints. Mol Biol Evol 20, 969–978. 10.1093/molbev/msg111.

4. Oggenfuss, U., Badet, T., Wicker, T., Hartmann, F.E., Singh, N.K., Abraham, L., Karisto, P., Vonlanthen, T., Mundt, C., McDonald, B.A., and Croll, D. (2021). A population-level invasion by transposable elements triggers genome expansion in a fungal pathogen. Elife 10. 10.7554/eLife.69249.

5. Gozashti, L., Roy, S.W., Thornlow, B., Kramer, A., Ares, M., Jr., and Corbett-Detig, R. (2022). Transposable elements drive intron gain in diverse eukaryotes. Proc Natl Acad Sci U S A 119, e2209766119. 10.1073/pnas.2209766119.

6. Lander, E.S., Linton, L.M., Birren, B., Nusbaum, C., Zody, M.C., Baldwin, J., Devon, K., Dewar, K., Doyle, M., FitzHugh, W., et al. (2001). Initial sequencing and analysis of the human genome. Nature 409, 860–921. 10.1038/35057062.

7. Etchegaray, E., Naville, M., Volff, J.N., and Haftek-Terreau, Z. (2021). Transposable element-derived sequences in vertebrate development. Mob DNA 12, 1. 10.1186/s13100-020-00229-5.

8. Wang, D., Su, Y., Wang, X., Lei, H., and Yu, J. (2012). Transposon-derived and satellite-derived repetitive sequences play distinct functional roles in Mammalian intron size expansion. Evol Bioinform Online 8, 301–319. 10.4137/EBO.S9758.

9. Zhang, X.H., Leslie, C.S., and Chasin, L.A. (2005). Computational searches for splicing signals. Methods 37, 292–305. 10.1016/j.ymeth.2005.07.011.

10. Cheng, Y., Kang, X.Z., Chan, P., Ye, Z.W., Chan, C.P., and Jin, D.Y. (2022). Aberrant splicing events caused by insertion of genes of interest into expression vectors. Int J Biol Sci 18, 4914–4931. 10.7150/ijbs.72408.

11. Black, D.L. (2005). A simple answer for a splicing conundrum. Proc Natl Acad Sci U S A 102, 4927–4928. 10.1073/pnas.0501414102.

12. Sun, H., and Chasin, L.A. (2000). Multiple splicing defects in an intronic false exon. Mol Cell Biol 20, 6414–6425. 10.1128/MCB.20.17.6414-6425.2000.

13. Humphrey, J., Emmett, W., Fratta, P., Isaacs, A.M., and Plagnol, V. (2017). Quantitative analysis of cryptic splicing associated with TDP-43 depletion. BMC Med Genomics 10, 38. 10.1186/s12920-017-0274-1.

14. Aldalaqan, S., Dalgliesh, C., Luzzi, S., Siachisumo, C., Reynard, L.N., Ehrmann, I., and Elliott, D.J. (2022). Cryptic splicing: common pathological mechanisms involved in male infertility and neuronal diseases. Cell Cycle 21, 219–227. 10.1080/15384101.2021.2015672.

15. Keegan, N.P., Wilton, S.D., and Fletcher, S. (2021). Analysis of Pathogenic Pseudoexons Reveals Novel Mechanisms Driving Cryptic Splicing. Front Genet 12, 806946. 10.3389/fgene.2021.806946.

16. Metherell, L.A., Akker, S.A., Munroe, P.B., Rose, S.J., Caulfield, M., Savage, M.O., Chew, S.L., and Clark, A.J. (2001). Pseudoexon activation as a novel mechanism for disease resulting in atypical growth-hormone insensitivity. Am J Hum Genet 69, 641–646. 10.1086/323266.

17. Glisovic, T., Bachorik, J.L., Yong, J., and Dreyfuss, G. (2008). RNA-binding proteins and post-transcriptional gene regulation. FEBS Lett 582, 1977–1986. 10.1016/j.febslet.2008.03.004.

18. Brinegar, A.E., and Cooper, T.A. (2016). Roles for RNA-binding proteins in development and disease. Brain Res 1647, 1–8. 10.1016/j.brainres.2016.02.050.

19. Qin, H., Ni, H., Liu, Y., Yuan, Y., Xi, T., Li, X., and Zheng, L. (2020). RNA-binding proteins in tumor progression. J Hematol Oncol 13, 90. 10.1186/s13045-020-00927-w.

20. Oliveira, C., Faoro, H., Alves, L.R., and Goldenberg, S. (2017). RNA-binding proteins and their role in the regulation of gene expression in Trypanosoma cruzi and Saccharomyces cerevisiae. Genet Mol Biol 40, 22–30. 10.1590/1678-4685-GMB-2016-0258.

21. Zarnack, K., Konig, J., Tajnik, M., Martincorena, I., Eustermann, S., Stevant, I., Reyes, A., Anders, S., Luscombe, N.M., and Ule, J. (2013). Direct competition between hnRNP C and U2AF65 protects the transcriptome from the exonization of Alu elements. Cell 152, 453–466. 10.1016/j.cell.2012.12.023.

22. Ma, X.R., Prudencio, M., Koike, Y., Vatsavayai, S.C., Kim, G., Harbinski, F., Briner, A., Rodriguez, C.M., Guo, C., Akiyama, T., et al. (2022). TDP-43 represses cryptic exon inclusion in the FTD-ALS gene UNC13A. Nature 603, 124–130. 10.1038/s41586-022-04424-7.

23. Attig, J., Agostini, F., Gooding, C., Chakrabarti, A.M., Singh, A., Haberman, N., Zagalak, J.A., Emmett, W., Smith, C.W.J., Luscombe, N.M., and Ule, J. (2018). Heteromeric RNP Assembly at LINEs Controls Lineage-Specific RNA Processing. Cell 174, 1067–1081 e1017. 10.1016/j.cell.2018.07.001.

24. Chen, Y.G., and Hur, S. (2022). Cellular origins of dsRNA, their recognition and consequences. Nat Rev Mol Cell Biol 23, 286–301. 10.1038/s41580-021-00430-1.

25. Hur, S. (2019). Double-Stranded RNA Sensors and Modulators in Innate Immunity. Annu Rev Immunol 37, 349–375. 10.1146/annurev-immunol-042718-041356.

26. Bowling, E.A., Wang, J.H., Gong, F., Wu, W., Neill, N.J., Kim, I.S., Tyagi, S., Orellana, M., Kurley, S.J., Dominguez-Vidana, R., et al. (2021). Spliceosome-targeted therapies trigger an antiviral immune response in triple-negative breast cancer. Cell 184, 384–403 e321. 10.1016/j.cell.2020.12.031.

27. Van Nostrand, E.L., Freese, P., Pratt, G.A., Wang, X., Wei, X., Xiao, R., Blue, S.M., Chen, J.Y., Cody, N.A.L., Dominguez, D., et al. (2020). A large-scale binding and functional map of human RNA-binding proteins. Nature 583, 711–719. 10.1038/s41586-020-2077-3.

28. Wang, T., Birsoy, K., Hughes, N.W., Krupczak, K.M., Post, Y., Wei, J.J., Lander, E.S., and Sabatini, D.M. (2015). Identification and characterization of essential genes in the human genome. Science 350, 1096–1101. 10.1126/science.aac7041.

29. Blomen, V.A., Majek, P., Jae, L.T., Bigenzahn, J.W., Nieuwenhuis, J., Staring, J., Sacco, R., van Diemen, F.R., Olk, N., Stukalov, A., et al. (2015). Gene essentiality and synthetic lethality in haploid human cells. Science 350, 1092–1096. 10.1126/science.aac7557.

30. Hart, T., Chandrashekhar, M., Aregger, M., Steinhart, Z., Brown, K.R., MacLeod, G., Mis, M., Zimmermann, M., Fradet-Turcotte, A., Sun, S., et al. (2015). High-Resolution CRISPR Screens Reveal Fitness Genes and Genotype-Specific Cancer Liabilities. Cell 163, 1515–1526. 10.1016/j.cell.2015.11.015.

31. Xu, Y., Gao, X.D., Lee, J.H., Huang, H., Tan, H., Ahn, J., Reinke, L.M., Peter, M.E., Feng, Y., Gius, D., et al. (2014). Cell type-restricted activity of hnRNPM promotes breast cancer metastasis via regulating alternative splicing. Genes Dev 28, 1191–1203. 10.1101/gad.241968.114.

32. Ramesh, N., Kour, S., Anderson, E.N., Rajasundaram, D., and Pandey, U.B. (2020). RNA-recognition motif in Matrin-3 mediates neurodegeneration through interaction with hnRNPM. Acta Neuropathol Commun 8, 138. 10.1186/s40478-020-01021-5.

33. Ho, J.S., Di Tullio, F., Schwarz, M., Low, D., Incarnato, D., Gay, F., Tabaglio, T., Zhang, J., Wollmann, H., Chen, L., et al. (2021). HNRNPM controls circRNA biogenesis and splicing fidelity to sustain cancer cell fitness. Elife 10. 10.7554/eLife.59654.

34. Wang, X., Li, J., Bian, X., Wu, C., Hua, J., Chang, S., Yu, T., Li, H., Li, Y., Hu, S., et al. (2021). CircURI1 interacts with hnRNPM to inhibit metastasis by modulating alternative splicing in gastric cancer. Proc Natl Acad Sci U S A 118. 10.1073/pnas.2012881118.

35. Shen, S., Park, J.W., Lu, Z.X., Lin, L., Henry, M.D., Wu, Y.N., Zhou, Q., and Xing, Y. (2014). rMATS: robust and flexible detection of differential alternative splicing from replicate RNA-Seq data. Proc Natl Acad Sci U S A 111, E5593–5601. 10.1073/pnas.1419161111.

36. Harvey, S.E., Xu, Y., Lin, X., Gao, X.D., Qiu, Y., Ahn, J., Xiao, X., and Cheng, C. (2018). Coregulation of alternative splicing by hnRNPM and ESRP1 during EMT. RNA 24, 1326–1338. 10.1261/rna.066712.118.

37. Huelga, S.C., Vu, A.Q., Arnold, J.D., Liang, T.Y., Liu, P.P., Yan, B.Y., Donohue, J.P., Shiue, L., Hoon, S., Brenner, S., et al. (2012). Integrative genome-wide analysis reveals cooperative regulation of alternative splicing by hnRNP proteins. Cell Rep 1, 167–178. 10.1016/j.celrep.2012.02.001.

38. Sibley, C.R., Blazquez, L., and Ule, J. (2016). Lessons from non-canonical splicing. Nat Rev Genet 17, 407–421. 10.1038/nrg.2016.46.

39. Smit, A.F. (1999). Interspersed repeats and other mementos of transposable elements in mammalian genomes. Curr Opin Genet Dev 9, 657–663. 10.1016/s0959-437x(99)00031-3.

40. Smit, A., Hubley, R., and Green, P. (1996-2004). RepeatMasker Open-3.0. http://wwwrepeatmasker.org.

41. Jaganathan, K., Kyriazopoulou Panagiotopoulou, S., McRae, J.F., Darbandi, S.F., Knowles, D., Li, Y.I., Kosmicki, J.A., Arbelaez, J., Cui, W., Schwartz, G.B., et al. (2019). Predicting Splicing from Primary Sequence with Deep Learning. Cell 176, 535–548 e524. 10.1016/j.cell.2018.12.015.

42. Cordaux, R., and Batzer, M.A. (2009). The impact of retrotransposons on human genome evolution. Nat Rev Genet 10, 691–703. 10.1038/nrg2640.

43. Lorenz, R., Bernhart, S.H., Honer Zu Siederdissen, C., Tafer, H., Flamm, C., Stadler, P.F., and Hofacker, I.L. (2011). ViennaRNA Package 2.0. Algorithms Mol Biol 6, 26. 10.1186/1748-7188-6-26.

44. Batzer, M.A., and Deininger, P.L. (2002). Alu repeats and human genomic diversity. Nat Rev Genet 3, 370–379. 10.1038/nrg798.

45. Walkley, C.R., and Li, J.B. (2017). Rewriting the transcriptome: adenosine-to-inosine RNA editing by ADARs. Genome Biol 18, 205. 10.1186/s13059-017-1347-3.

46. Hong, H., Lin, J.S., and Chen, L. (2015). Regulatory factors governing adenosine-to-inosine (A-to-I) RNA editing. Biosci Rep 35. 10.1042/BSR20140190.

47. Wang, Q., and Carmichael, G.G. (2004). Effects of length and location on the cellular response to double-stranded RNA. Microbiol Mol Biol Rev 68, 432–452, table of contents. 10.1128/MMBR.68.3.432-452.2004.

48. Weber, F., Wagner, V., Rasmussen, S.B., Hartmann, R., and Paludan, S.R. (2006). Double-stranded RNA is produced by positive-strand RNA viruses and DNA viruses but not in detectable amounts by negative-strand RNA viruses. J Virol 80, 5059–5064. 10.1128/JVI.80.10.5059-5064.2006.

49. Werner, A., Clark, J.E., Samaranayake, C., Casement, J., Zinad, H.S., Sadeq, S., Al-Hashimi, S., Smith, M., Kotaja, N., and Mattick, J.S. (2021). Widespread formation of double-stranded RNAs in testis. Genome Res 31, 1174–1186. 10.1101/gr.265603.120.

50. Onomoto, K., Onoguchi, K., and Yoneyama, M. (2021). Regulation of RIG-I-like receptor-mediated signaling: interaction between host and viral factors. Cell Mol Immunol 18, 539–555. 10.1038/s41423-020-00602-7.

51. Qiao, L., Xie, N., Li, Y., Bai, Y., Liu, N., and Wang, J. (2022). Downregulation of HNRNPM inhibits cell proliferation and migration of hepatocellular carcinoma through MAPK/AKT signaling pathway. Transl Cancer Res 11, 2135–2144. 10.21037/tcr-21-2484.

52. Vitali, P., and Scadden, A.D. (2010). Double-stranded RNAs containing multiple IU pairs are sufficient to suppress interferon induction and apoptosis. Nat Struct Mol Biol 17, 1043–1050. 10.1038/nsmb.1864.

53. Liberzon, A., Birger, C., Thorvaldsdottir, H., Ghandi, M., Mesirov, J.P., and Tamayo, P. (2015). The Molecular Signatures Database (MSigDB) hallmark gene set collection. Cell Syst 1, 417–425. 10.1016/j.cels.2015.12.004.

54. Arico, E., Castiello, L., Capone, I., Gabriele, L., and Belardelli, F. (2019). Type I Interferons and Cancer: An Evolving Story Demanding Novel Clinical Applications. Cancers (Basel) 11. 10.3390/cancers11121943.

55. Cao, X., Liang, Y., Hu, Z., Li, H., Yang, J., Hsu, E.J., Zhu, J., Zhou, J., and Fu, Y.X. (2021). Next generation of tumor-activating type I IFN enhances anti-tumor immune responses to overcome therapy resistance. Nat Commun 12, 5866. 10.1038/s41467-021-26112-2.

56. Zitvogel, L., Galluzzi, L., Kepp, O., Smyth, M.J., and Kroemer, G. (2015). Type I interferons in anticancer immunity. Nat Rev Immunol 15, 405–414. 10.1038/nri3845.

57. Yi, M., Nissley, D.V., McCormick, F., and Stephens, R.M. (2020). ssGSEA score-based Ras dependency indexes derived from gene expression data reveal potential Ras addiction mechanisms with possible clinical implications. Sci Rep 10, 10258. 10.1038/s41598-020-66986-8.

58. Kramer, B., Knoll, R., Bonaguro, L., ToVinh, M., Raabe, J., Astaburuaga-Garcia, R., Schulte-Schrepping, J., Kaiser, K.M., Rieke, G.J., Bischoff, J., et al. (2021). Early IFN-alpha signatures and persistent dysfunction are distinguishing features of NK cells in severe COVID-19. Immunity 54, 2650–2669 e2614. 10.1016/j.immuni.2021.09.002.

59. Fenton, S.E., Saleiro, D., and Platanias, L.C. (2021). Type I and II Interferons in the Anti-Tumor Immune Response. Cancers (Basel) 13. 10.3390/cancers13051037.

60. Crouse, J., Kalinke, U., and Oxenius, A. (2015). Regulation of antiviral T cell responses by type I interferons. Nat Rev Immunol 15, 231–242. 10.1038/nri3806.

61. Aran, D., Hu, Z., and Butte, A.J. (2017). xCell: digitally portraying the tissue cellular heterogeneity landscape. Genome Biol 18, 220. 10.1186/s13059-017-1349-1.

62. Geuens, T., Bouhy, D., and Timmerman, V. (2016). The hnRNP family: insights into their role in health and disease. Hum Genet 135, 851–867. 10.1007/s00439-016-1683-5.

63. Hovhannisyan, R.H., and Carstens, R.P. (2007). Heterogeneous ribonucleoprotein m is a splicing regulatory protein that can enhance or silence splicing of alternatively spliced exons. J Biol Chem 282, 36265–36274. 10.1074/jbc.M704188200.

64. Garcia-Garijo, A., Fajardo, C.A., and Gros, A. (2019). Determinants for Neoantigen Identification. Front Immunol 10, 1392. 10.3389/fimmu.2019.01392.

65. Kahles, A., Lehmann, K.V., Toussaint, N.C., Huser, M., Stark, S.G., Sachsenberg, T., Stegle, O., Kohlbacher, O., Sander, C., Cancer Genome Atlas Research, N., and Ratsch, G. (2018). Comprehensive Analysis of Alternative Splicing Across Tumors from 8,705 Patients. Cancer Cell 34, 211–224 e216. 10.1016/j.ccell.2018.07.001.

66. Seddighi, S., Qi, Y.A., Brown, A.L., Wilkins, O.G., Bereda, C., Belair, C., Zhang, Y., Prudencio, M., Keuss, M.J., Khandeshi, A., et al. (2023). Mis-spliced transcripts generate de novo proteins in TDP-43-related ALS/FTD. bioRxiv. 10.1101/2023.01.23.525149.

67. West, K.O., Scott, H.M., Torres-Odio, S., West, A.P., Patrick, K.L., and Watson, R.O. (2019). The Splicing Factor hnRNP M Is a Critical Regulator of Innate Immune Gene Expression in Macrophages. Cell Rep 29, 1594–1609 e1595. 10.1016/j.celrep.2019.09.078.

68. Cao, P., Luo, W.W., Li, C., Tong, Z., Zheng, Z.Q., Zhou, L., Xiong, Y., and Li, S. (2019). The heterogeneous nuclear ribonucleoprotein hnRNPM inhibits RNA virus-triggered innate immunity by antagonizing RNA sensing of RIG-I-like receptors. PLoS Pathog 15, e1007983. 10.1371/journal.ppat.1007983.

69. Choi, H., Kwon, J., Cho, M.S., Sun, Y., Zheng, X., Wang, J., Bouker, K.B., Casey, J.L., Atkins, M.B., Toretsky, J., and Han, C. (2021). Targeting DDX3X Triggers Antitumor Immunity via a dsRNA-Mediated Tumor-Intrinsic Type I Interferon Response. Cancer Res 81, 3607–3620. 10.1158/0008-5472.CAN-20-3790.

70. Shi, C., Wang, Y., Wu, M., Chen, Y., Liu, F., Shen, Z., Wang, Y., Xie, S., Shen, Y., Sang, L., et al. (2022). Promoting anti-tumor immunity by targeting TMUB1 to modulate PD-L1 polyubiquitination and glycosylation. Nat Commun 13, 6951. 10.1038/s41467-022-34346-x.

71. Hong, S. (2017). RNA Binding Protein as an Emerging Therapeutic Target for Cancer Prevention and Treatment. J Cancer Prev 22, 203–210. 10.15430/JCP.2017.22.4.203.

72. Mohibi, S., Chen, X., and Zhang, J. (2019). Cancer the’RBP’eutics-RNA-binding proteins as therapeutic targets for cancer. Pharmacol Ther 203, 107390. 10.1016/j.pharmthera.2019.07.001.

73. Kang, D., Lee, Y., and Lee, J.S. (2020). RNA-Binding Proteins in Cancer: Functional and Therapeutic Perspectives. Cancers (Basel) 12. 10.3390/cancers12092699.

74. Damianov, A., Ying, Y., Lin, C.H., Lee, J.A., Tran, D., Vashisht, A.A., Bahrami-Samani, E., Xing, Y., Martin, K.C., Wohlschlegel, J.A., and Black, D.L. (2016). Rbfox Proteins Regulate Splicing as Part of a Large Multiprotein Complex LASR. Cell 165, 606–619. 10.1016/j.cell.2016.03.040.

75. Dobin, A., Davis, C.A., Schlesinger, F., Drenkow, J., Zaleski, C., Jha, S., Batut, P., Chaisson, M., and Gingeras, T.R. (2013). STAR: ultrafast universal RNA-seq aligner. Bioinformatics 29, 15–21. 10.1093/bioinformatics/bts635.

76. Love, M.I., Huber, W., and Anders, S. (2014). Moderated estimation of fold change and dispersion for RNA-seq data with DESeq2. Genome Biol 15, 550. 10.1186/s13059-014-0550-8.

77. Subramanian, A., Tamayo, P., Mootha, V.K., Mukherjee, S., Ebert, B.L., Gillette, M.A., Paulovich, A., Pomeroy, S.L., Golub, T.R., Lander, E.S., and Mesirov, J.P. (2005). Gene set enrichment analysis: a knowledge-based approach for interpreting genome-wide expression profiles. Proc Natl Acad Sci U S A 102, 15545–15550. 10.1073/pnas.0506580102.

78. Liao, Y., Wang, J., Jaehnig, E.J., Shi, Z., and Zhang, B. (2019). WebGestalt 2019: gene set analysis toolkit with revamped UIs and APIs. Nucleic Acids Res 47, W199–W205. 10.1093/nar/gkz401.

79. Harvey, S.E., and Cheng, C. (2016). Methods for Characterization of Alternative RNA Splicing. Methods Mol Biol 1402, 229–241. 10.1007/978-1-4939-3378-5_18.

80. Quinones-Valdez, G., Tran, S.S., Jun, H.I., Bahn, J.H., Yang, E.W., Zhan, L., Brummer, A., Wei, X., Van Nostrand, E.L., Pratt, G.A., et al. (2019). Regulation of RNA editing by RNA-binding proteins in human cells. Commun Biol 2, 19. 10.1038/s42003-018-0271-8.

81. Bahn, J.H., Lee, J.H., Li, G., Greer, C., Peng, G., and Xiao, X. (2012). Accurate identification of A-to-I RNA editing in human by transcriptome sequencing. Genome Res 22, 142–150. 10.1101/gr.124107.111.

82. Lee, J.H., Ang, J.K., and Xiao, X. (2013). Analysis and design of RNA sequencing experiments for identifying RNA editing and other single-nucleotide variants. RNA 19, 725–732. 10.1261/rna.037903.112.

83. Zhang, Q., and Xiao, X. (2015). Genome sequence-independent identification of RNA editing sites. Nat Methods 12, 347–350. 10.1038/nmeth.3314.

84. Yang, E.W., Bahn, J.H., Hsiao, E.Y., Tan, B.X., Sun, Y., Fu, T., Zhou, B., Van Nostrand, E.L., Pratt, G.A., Freese, P., et al. (2019). Allele-specific binding of RNA-binding proteins reveals functional genetic variants in the RNA. Nat Commun 10, 1338. 10.1038/s41467-019-09292-w.

